# Two alveolin network proteins are essential for the subpellicular microtubules assembly and conoid anchoring to the apical pole of mature *Toxoplasma gondii*

**DOI:** 10.1101/2020.02.25.962589

**Authors:** Nicolò Tosetti, Nicolas Dos Santos Pacheco, Eloïse Bertiaux, Bohumil Maco, Lorène Bournonville, Virginie Hamel, Paul Guichard, Dominique Soldati-Favre

## Abstract

*Toxoplasma gondii* belongs to the coccidian sub-group of Apicomplexa that possess an apical complex harboring a conoid, made of unique tubulin polymer fibers. This enigmatic and dynamic organelle extrudes in extracellular invasive parasites and is associated to the apical polar ring (APR), a microtubule-organizing center for the 22 subpellicular microtubules (SPMTs). The SPMTs are linked to the Inner Membrane Complex (IMC), a patchwork of flattened vesicles, via an intricate network of small filaments composed of alveolins proteins. Here, we capitalize on super-resolution techniques including stimulated emission depletion (STED) microscopy and ultrastructure expansion microscopy (U-ExM) to localize the Apical Cap protein 9 (AC9) and its close partner AC10, identified by BioID, to the alveolin network and intercalated between the SPMTs. Conditional depletion of AC9 or AC10 using the Auxin-induced Degron (AiD) system uncovered a severe loss of fitness. Parasites lacking AC9 or AC10 replicate normally but are defective in microneme secretion and hence fail to invade and egress from infected cells. Remarkably, a series of crucial apical complex proteins (MyoH, AKMT, FRM1, CPH1, ICMAP1 and RNG2) are lost in the mature parasites although they are still present in the forming daughter cells. Electron microscopy on intracellular or deoxycholate-extracted parasites revealed that the mature parasite mutants are conoidless. Closer examination of the SPMTs by U-ExM highlighted the disassembly of the SPMTs in the apical cap region that is presumably at the origin of the catastrophic loss of APR and conoid. AC9 and AC10 are two critical components of the alveolin network that ensure the integrity of the whole apical complex in *T. gondii* and likely other coccidians.

## Introduction

*Toxoplasma gondii* belongs to the phylum of Apicomplexa that groups numerous parasitic protozoans causing severe diseases in humans and animals. As part of the superphylum of Alveolata, the Apicomplexa are characterized by the presence of the alveoli, which consist in small flattened single-membrane sacs, underlying the plasma membrane (PM) to form the inner membrane complex (IMC) of the parasite. A rigid alveolin network composed of intermediate filament-like proteins is lining the cytoplasmic side of the IMC. The alveolin network is made of proteins sharing conserved repeat motifs called alveolins, which together with the IMC span the length of the parasite from the apical polar ring (APR) to the basal complex (1-3). The IMC plays an essential role in parasite motility by anchoring the actomyosin system (4) and serves as structural scaffold during daughter cells formation within the mother cell, an asexual form of reproduction referred as endodyogeny (5).

The IMC is arranged in a series of rectangular plates sutured together. Several IMC sutures components (ISCs) have been localized to the transversal and longitudinal sutures between alveolar sacs (6, 7). The parasite is capped by a single cone-shaped plate called the apical cap. Several proteins have been reported to localize at the apical cap including the IMC sub-compartment protein 1 (ISP1) (8) and nine apical cap proteins (called AC1 to AC9) (6, 7). Centrin2 labels a peripheral ring of six annuli at the boundary of the apical plate and the rest of the alveolar plates (9). Beneath the alveolin network, a set of 22 subpellicular microtubules (SPMT) spanning two third of the parasite length confer the elongated shape to the tachyzoites. The SPTMs originate at the APR, which serves as microtubule-organizing center (MTOC). Ultrastructural studies have highlighted tight connections between the IMC complex and the SPMTs. More specifically, freeze-fractured studies showed double rows of inner membranous particles (IMPs) at the IMC surface arranged in spiraling longitudinal rows reminiscent of SPMT path exhibiting a repeating pattern with a 32 nm periodicity. This 32 nm periodicity was also observed in the single row of IMP associated with SPMTs (10).

The APR and the secretory organelles, rhoptries (implicated in invasion) and micronemes (implicated in motility, invasion and egress), are common features of the apical complex conserved in all motile and invasive apicomplexans. Members of the coccidian subgroup of the phylum, including *Toxoplasma, Neospora, Besniotia, Cyclospora, Sarcocystis* and *Eimeria*, possess an additional organelle termed the conoid. This dynamic organelle is formed by 14 atypical comma-shape tubulin fibers composed of ∼9 protofilaments differing from classical microtubules (11). The cone of the conoid is enclosed between the APR and two preconoidal rings (PCRs) at the apex of the parasite. In addition, two short intraconoidal microtubules are present within the conoid (12). During division, the conoid is retracted posterior to the APR whereas it protrudes through the APR in a calcium-dependent manner in motile parasites (13). The calcium-dependent protein kinase 1 (CDPK1) is an essential regulator of microneme exocytosis (14) and its deletion results in inhibition of conoid protrusion and a block of the apico-basal flux of F-actin, essential for parasite motility (15). The mechanistic contribution of the conoid in motility and invasion has not been elucidated to date. During evolution, the conoid was lost in Haematozoa which include *Plasmodium* spp. and piroplasmidia (*Theileria* and *Babesia*). In contrast, the conoid is still present in deeply branching *Cryptosporidium* spp. and *Gregarina* spp. Other more distantly related Alveolata (colpodellids, perkinsids and chromerids), possess a similar structure referred to as incomplete conoid or pseudoconoid, built from apical MTs but lacking the APR, which suggests that ancestral apicomplexans harbored an ancestral apical complex along with secretory organelles (16). Intriguingly, the SAS6-like protein, which is localized near the base of the flagellum of *Trypanosoma brucei*, is found at the preconoidal rings of *T. gondii* tachyzoite that do not possess a flagellum, raising questions about the origin of the conoid (17). SAS6-like has also been localized apically in both ookinetes and sporozoites but absent in merozoites of *Plasmodium berghei*, revealing that the morphology and composition of the apical complex might vary between different parasitic life stages and yet retained conserved proteins (18).

More than 250 proteins were enriched in the conoid fraction by a proteomic approach (9) but only a few have been functionally investigated. Knockdown of the APR protein RNG2 results in microneme secretion defect (19), while the conoid-associated MyoH and the likely associated calmodulin-like proteins are all required for gliding motility that powers invasion and egress (20, 21). Additionally, the apical complex methyltransferase (AKMT) (22), the formin 1 (FRM1) (15) and the glideosome-associated connector (GAC) (23) are apical proteins conserved across the Apicomplexa that play essential roles in parasite motility, invasion and egress (4). Among the proteins involved in stability of the apical complex, loss of microtubules binding doublecortin (DCX)-domain protein results in shorter disordered conoid with subsequent defect in invasion (24). DCX was later shown to bundle the conoidal tubulin fibers into comma-shape structures likely conferring the cone-shaped configuration of the conoid (25). Moreover, the double knockout of kinesin A and apical polar ring protein 1 (APR1) caused a fragmentation of the APR with a conoid frequently partially detached resulting in impairment of microneme secretion (26). Lastly, conoid protein hub 1 (CPH1) is a conserved apicomplexan protein essential for parasite motility, invasion and egress that regulates conoid stability in extracellular parasites without impacting on microneme secretion (27). Similarly, a recently described protein, the MAP kinase ERK7, localizing to the apical cap was proposed to be involved in conoid assembly without impacting on microneme secretion (28). Here, we have used ultrastructure expansion microscopy (U-ExM) to precisely localize AC9 and its partner AC10, identified by BioID, to the alveolin network but exclusively in the apical cap region in-between the SPMTs. Conditional depletion of AC9 and AC10 established that the two proteins are essential for parasite motility invasion and egress, while parasite replication is not affected. U-ExM and electron microscopy revealed striking morphological defects. AC9 or AC10 depleted mature parasites are conoidless, deprived of APR and the SPMTs are disassembled. The arrangement of AC9 and AC10 suggests that both protein act as a glue between SPMTs, a prerequisite to maintain the overall architecture of the key components of the apical complex when the parasite mature.

## Results

### AC10 is a new apical cap protein closely associated to AC9

*T. gondii* AC9 is an hypothetical protein exhibiting no informative protein domain to unravel its function but a strong negative fitness score indicative of essentiality (29). AC9 was modified by CRISPR/Cas9 editing at the endogenous locus to generate a fusion with the mini auxin-induced degron (mAID) and HA-tag at the C-terminus to localize and rapidly downregulate the protein upon addition of auxin (IAA) (30) (S1A Fig). As previously reported (7), AC9 colocalizes with ISP1 at the most apical plate of the IMC termed apical cap, delimited by the peripheral annuli stained by centrin2 (Cen2). Remarkably, AC9 appears very early during endodyogeny before the emergence of the daughter parasite scaffold stained with IMC1 antibodies (Fig 1A and S1B Fig). Further colocalization with other Ty-tagged proteins at their endogenous loci, revealed that AC9 surrounds the APR marked by RNG2-Ty (19) and the conoid localized with CPH1-Ty (Fig 1B). A second copy of the gliding associated protein 70 (GAP70)-mycGFP was introduced in the AC9-mAID-HA strain. Like GAP45, GAP70 is acylated and anchored at both the IMC and the PM spanning the space between those two membranes (31). GAP70-mycGFP localized slightly more apically compared to AC9, which remains confined to the apical plate of the IMC (Fig 1C). Next, fractionation assay revealed that AC9 is completely resistant to extraction with the non-ionic detergent Triton X-100 but is solubilized in sodium carbonate at pH 11. This suggests that AC9 is an alveolin network resident protein interacting via protein-protein interactions with other IMC/alveolins proteins (Fig 1D). Noteworthy, AC9-mAID-HA migrated on SDS-PAGE at around 70 kDa, slightly higher than the predicted size (60 kDa). To get insights into AC9 potential interacting partners, we opted for the proximity labeling by the biotin ligase BirA approach (32) given the limited solubility of AC9. A mycBirA construct was inserted at the endogenous C-terminus of AC9 by CRISPR/Cas9 (S1C Fig) and shown to be correctly targeted to the apical cap (S1D Fig). To assess whether AC9-mycBirA was able to biotinylate proximal proteins, we performed IFA with fluorophore-conjugated streptavidin. In absence of biotin, apicoplast endogenously biotinylated proteins were observed while addition of biotin for 24h resulted in accumulation of biotinylated proteins in the apical cap, proving that the enzyme is active at the right position (S1D Fig). Of relevance, deoxycholate extracted parasites showed a robust streptavidin staining around the apical cortical microtubules just beneath the conoid indicating biotinylation of components of the parasite cytoskeleton (S1D Fig). Analysis of parasites total lysates by WB showed an enrichment of biotinylated proteins (S1E Fig). Immunoprecipitation with streptavidin magnetic beads was performed with parasites lysed in cytoskeletal buffer as previously described (32) (S1F Fig) followed by mass spectrometry analysis. The clear top hit was a hypothetical protein (TGGT1_292950) followed by two apical cap proteins (AC2 and AC8) and IMC/cytoskeletal proteins (S1 Table). Candidates were further filtered by expression patterns similar to AC9 (S1G Fig). TGGT1_292950 exhibited a very similar cyclic mRNA expression pattern compared to AC9 (Fig 1E). Upon C-terminal Ty-epitope tagging, the product of the gene colocalized with AC9 in mature parasites and daughter cells and hence was named AC10 (Fig 1F). Like AC9, AC10 was showed to be partially solubilized by sodium carbonate and completely insoluble in Triton X-100 using the AC10-mAID-HA strain, indicating its alveolin network association (Fig 1G).

**Figure 1.**
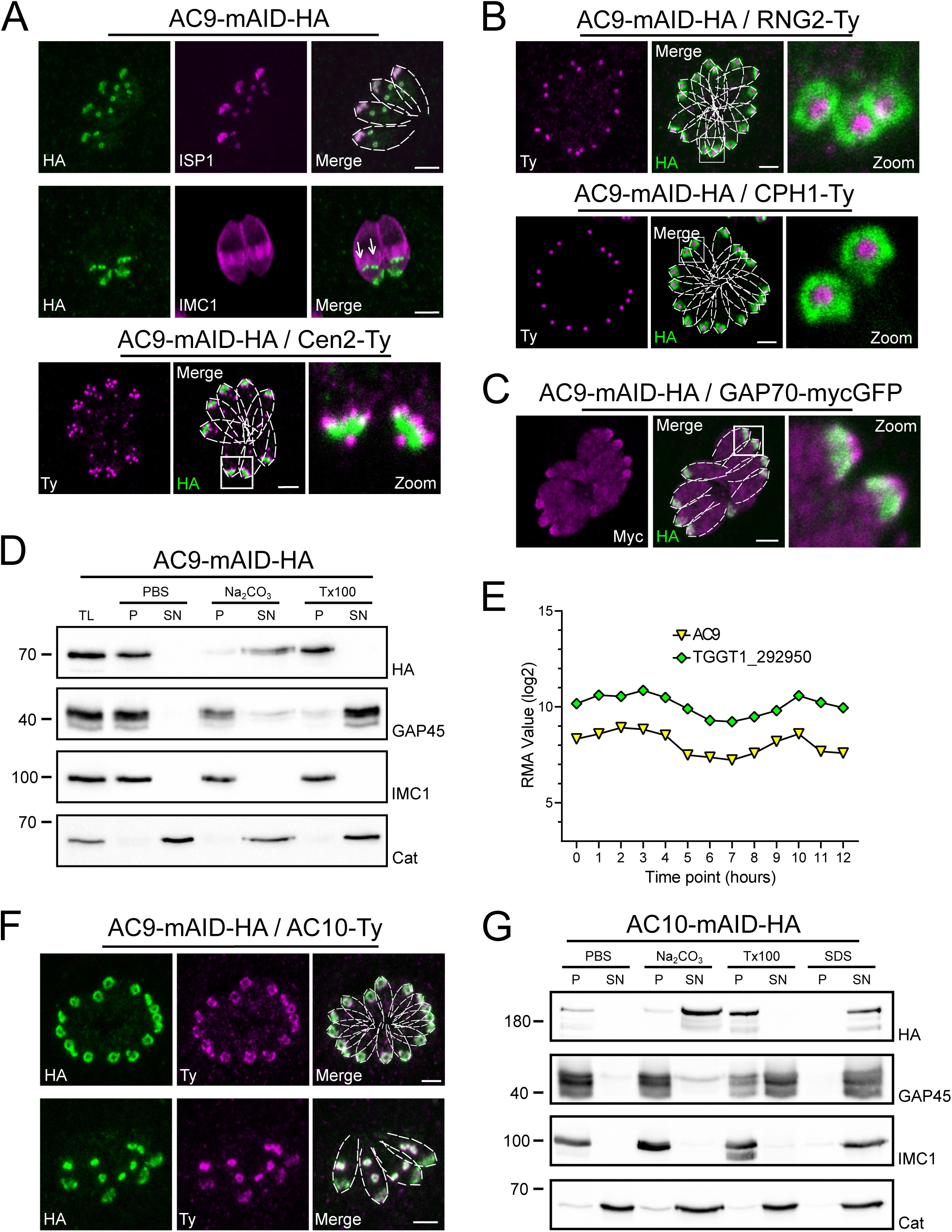
AC9 and the newly discovered AC10 are alveolin network proteins. **(A)** AC9-mAID-HA localized to the apical cap of both mature and daughters cells (arrows). AC9 is confined to the apical IMC plate delimited by the Centrin2 marker. **(B)** AC9 surrounds the conoid (CPH1 marker) and the apical polar ring (RNG2 marker). **(C)** Colocalization with GAP70 revealed that AC9 is likely confined in the IMC/subpellicular network (SPN) and not associated with the PM. **(D)** AC9-mAID-HA is partially solubilized by sodium carbonate and precipitated in Triton X-100 insoluble fraction. **(E)** TGGT1_292950 was the top biotinylated hit. This hypothetical protein displayed an identical cyclic expression pattern compared to AC9. **(F)** TGGT1_292950 colocalized with AC9 in both daughter and mature parasite and was subsequently named AC10. **(G)** Similar to AC9, AC10-mAID-HA is partially solubilized only by sodium carbonate. Scale bars = 2µm.

### AC9 and AC10 are recruited very early during daughter cells formation

ISP1 was localized to the apical cap along with several apical cap proteins (ACs) (6-8). Intriguingly, some ACs are recruited early during daughter cells formation while other appear later, highlighting the existence of a temporal and spatial hierarchy of IMC and alveolin network formation during parasite endodyogeny (6). Colocalization experiments revealed that AC9 and AC10 are recruited very early to the daughter cytoskeleton, prior to ISP1 incorporation (Fig 2A and 2B); in contrast, AC2 and AC8 are recruited later (S2A and S2B Fig), as schematized in Fig 2C. The three heavily biotinylated proteins AC2, AC8 and AC10, were endogenously tagged at the C-terminus in the AC9-mAID-HA strain to assess further the interaction with AC9. Deletion of AC9 had no impact on the localization of AC2, AC8, AC10 and ISP1 (Fig 2D and S2A-C Fig). Conversely, epitope tagged AC9 in AC10-mAID-HA disappeared upon AC10 depletion both by western blot (WB) and indirect immunofluorescence assay (IFA) (Fig 2E and 2F) providing additional compelling evidence that the two proteins form a complex. Even a short treatment with IAA was sufficient to cause destabilization of AC9 (Fig 2G) without impacting on AC2, AC8 and ISP1 (S2D-F Fig), yet the apical pole appeared enlarged (arrow) in some parasites (S2C and S2F Fig).

**Figure 2.**
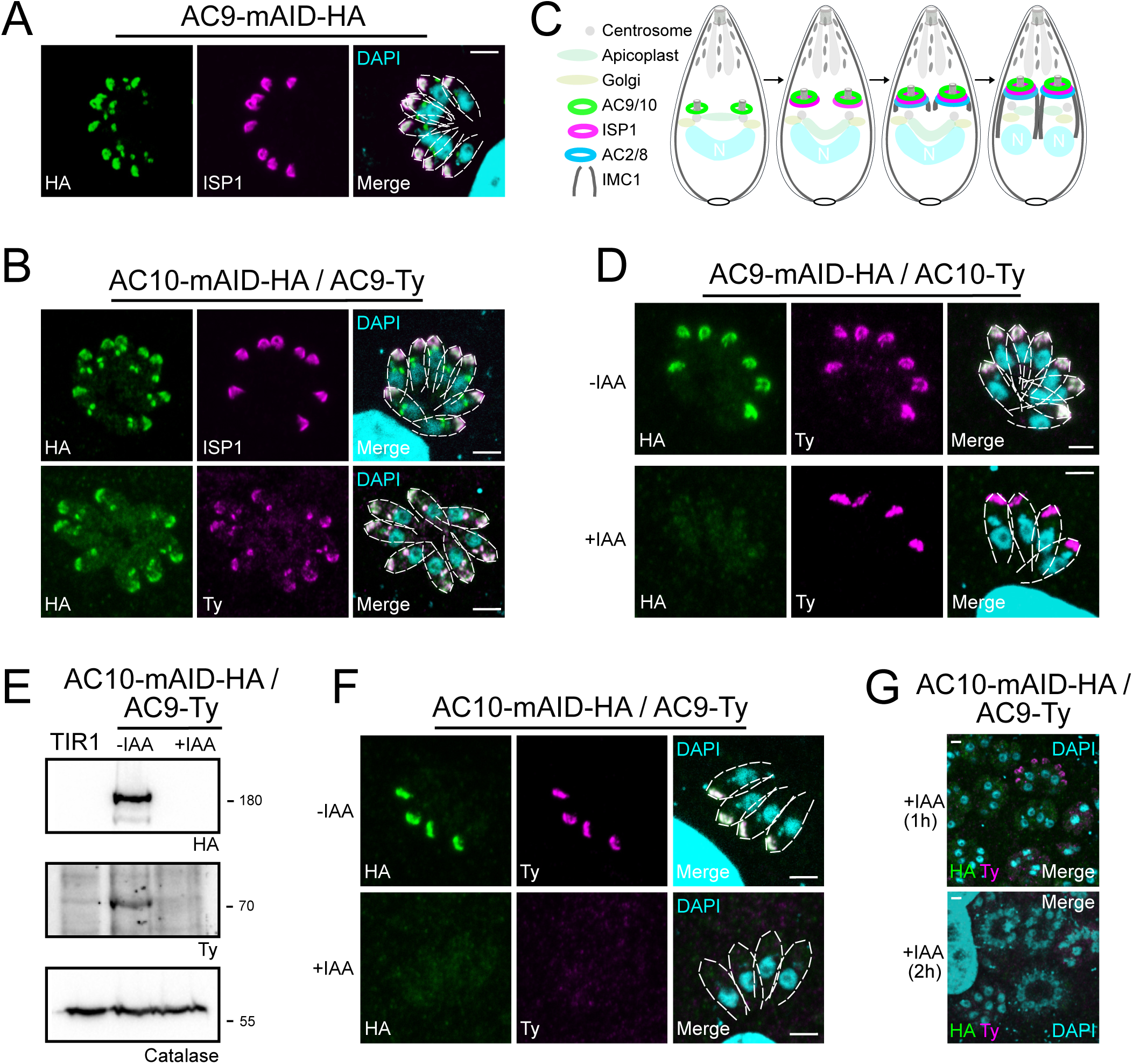
AC9 and AC10 are among the earliest markers of parasite division. **(A)** and **(B)** Both AC9 and AC10 are both recruited very early during budding of daughters cells and before ISP1. **(C)** Schematic representation of the events during endodyogeny highlighting the sequential insertion of the different components of the IMC/SPN/cytoskeleton. **(D)** AC10 localization was not impacted upon depletion of AC9. **(E)** and **(F)** Depletion of AC10 caused degradation of AC9. **(G)** Shorter auxin treatments were sufficient to remove AC9 from the apical cap suggesting that AC9 and AC10 forms a complex. Scale bars = 2µm.

### AC9 and AC10 belong to the alveolin network based on super-resolution microscopy

Super-resolution imaging with a stimulated emission depletion (STED) microscope revealed that AC9 and AC10 signals are not homogeneous at the apical cap but rather organized in rows with a regular periodicity (Fig 3A and 3B). Remarkably, IMC1 staining is also organized in regular rows reminiscent of the SPMTs arrangement and the intramembranous particle lattice observed by electron microscopy (EM) (10). Further colocalization of IMC1 with tubulin showed that microtubules are interspaced by two rows of IMC1 (S3A Fig). In contrast, GAP45 staining is more homogeneous along parasite pellicle on the PM side of the IMC. Colocalization with GAP45 and IMC sub-compartment protein 1 (ISP1) (8) confirmed that AC9 is confined to the SPMTs side of the IMC and follows a periodic arrangement like SPMTs and alveolins (S3B Fig). We next applied Ultrastructure Expansion Microscopy (U-ExM) (33) for the first time in *T. gondii* to gain further resolution. As proof of concept, we first used antibodies specific to alpha/beta tubulin and to poly-glutamylation (Poly-E) to detect the SPMTs. Remarkably, the shape and ultrastructure of the parasite were preserved while the expansion rate was approaching 4x and the SPMTs showed a high level of poly-glutamylation all along their length, except at their most distal part (Fig S3C). Interestingly the conoid fibers seem to be devoid of poly-glutamylation. Of relevance, AC9 and AC10 were clearly colocalizing as a regular pattern between each SPMTs just below the conoid and APR (Fig 3C-E and S3D Fig). Given the close proximity to the SPMTs, AC9 and AC10 were produced recombinantly to assess binding to MTs in an *in vitro* binding assay (Cytoskeleton ink.). AC9 did not interact directly with MTs (S3E Fig) while recombinant AC10 produced either in bacteria or insect cells formed a gel resistant to solubilization and hence could not be tested. Analysis of the other ACs biotinylated proteins by AC9 revealed that AC2 is localized on the SPMTs up to the APR in contrast to AC8, which is present between the SPMTs and does not reach the APR (Fig 3F and 3G). STED microscopy confirmed that AC9 and AC10 are surrounding the APR stained by RNG2 without being in direct contact with it (Fig 3F). Of interest, RNG2 adopted a regular pattern at the APR both by STED and U-ExM, suggesting that the APR is composed of discrete subunits possibly including microtubule plus-end tracking proteins that ensure the apical docking of the SPMTs and their regular interspacing (Fig 3H and 3I).

**Figure 3.**
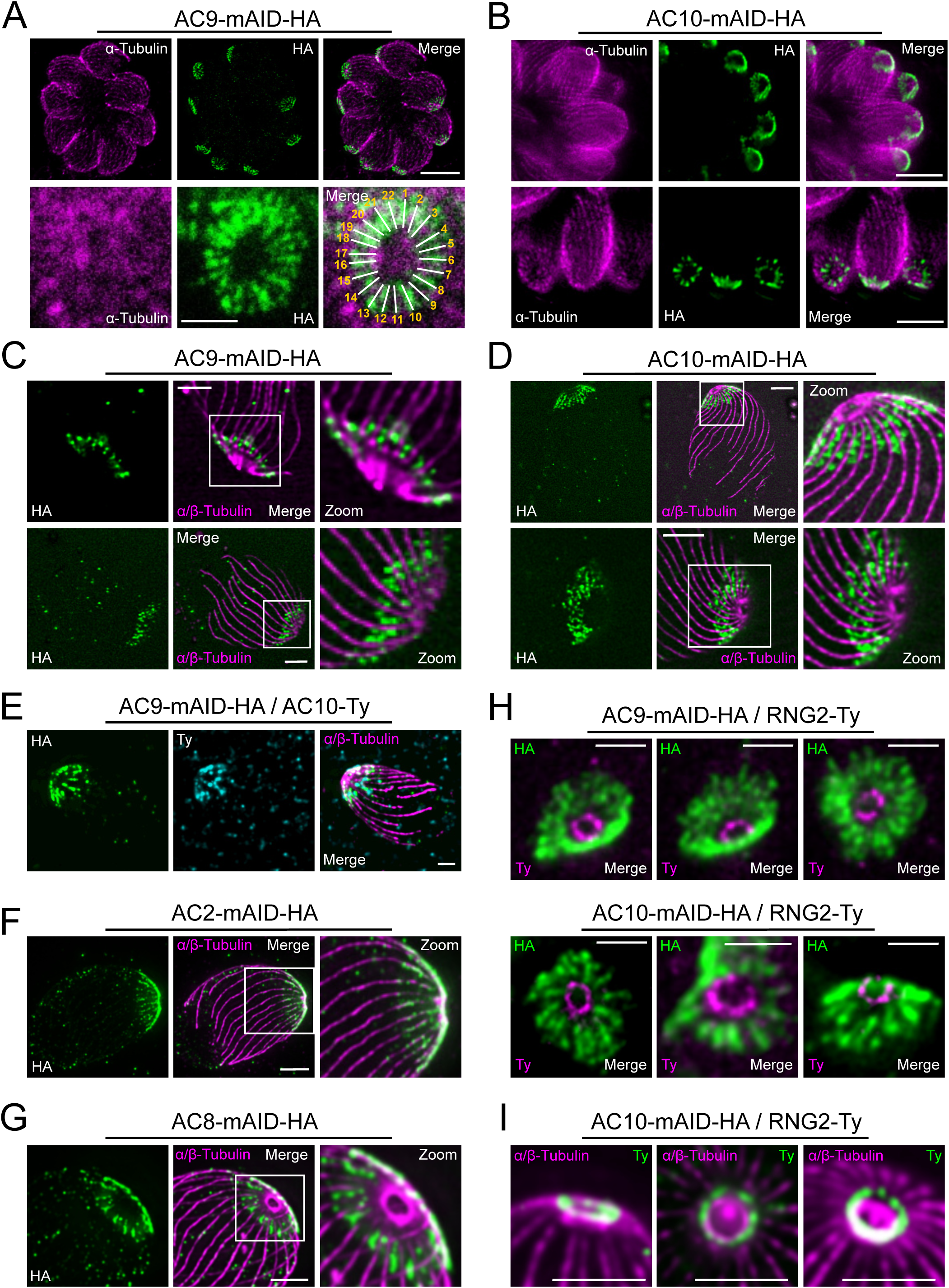
Super resolution microscopy revealed a peculiar localization of ACs proteins. **(A)** and **(B)** Stimulated emission-depletion super-resolution microscopy (STED) revealed that AC9 and AC10 is arranged in periodical rows reminiscent of subpellicular microtubules (SPMTs) arrangement. **(C)** and **(D)** Expansion microscopy (U-ExM) confirmed that that AC9 and AC10 are arranged in regular rows and localized in between the SPMTs. **(E)** AC9 and AC10 colocalized at the apical cap level. **(F)** AC2 localized on SPMTs by U-ExM at the apical cap up to the APR. **(G)** As AC9 and AC10, AC8 localized in between SPMTs. **(H)** and **(I)** Colocalization with RNG2 revealed that both AC9 and AC10 surrounds the APR without making significant contacts with it. Surprisingly, the RNG2 signal did not appear to be homogeneous at the APR but rather forming discrete dots by STED and U-ExM. Scale bars = 2µm, except for 1A (bottom panel) and 1H = 0.5 µm.

### AC9 and AC10 are essential for invasion and egress and for induced microneme secretion

Upon addition of auxin, AC9 and AC10 are tightly downregulated as shown by IFA (Fig 4A and 4D) and WB (Fig 4B and 4E) in the AC9-mAID-HA and AC10-mAID-HA strains. Depletion of both proteins resulted in no lysis plaques in the monolayer of human foreskin fibroblasts (HFF) after 7 days (Fig 4C and 4F). Further phenotyping showed that depletion of AC9 and AC10 caused a severe defect in invasion (Fig 4G) and egress (Fig 4H) without impacting on parasite intracellular growth and replication (S4A Fig). This phenotype is at least in part explained by the block in induced microneme secretion observed upon stimulation by two known triggers of microneme exocytosis, BIPPO (a phosphodiesterase inhibitor) (34) and ethanol (35) (Fig 4I and S4B Fig).

**Figure 4.**
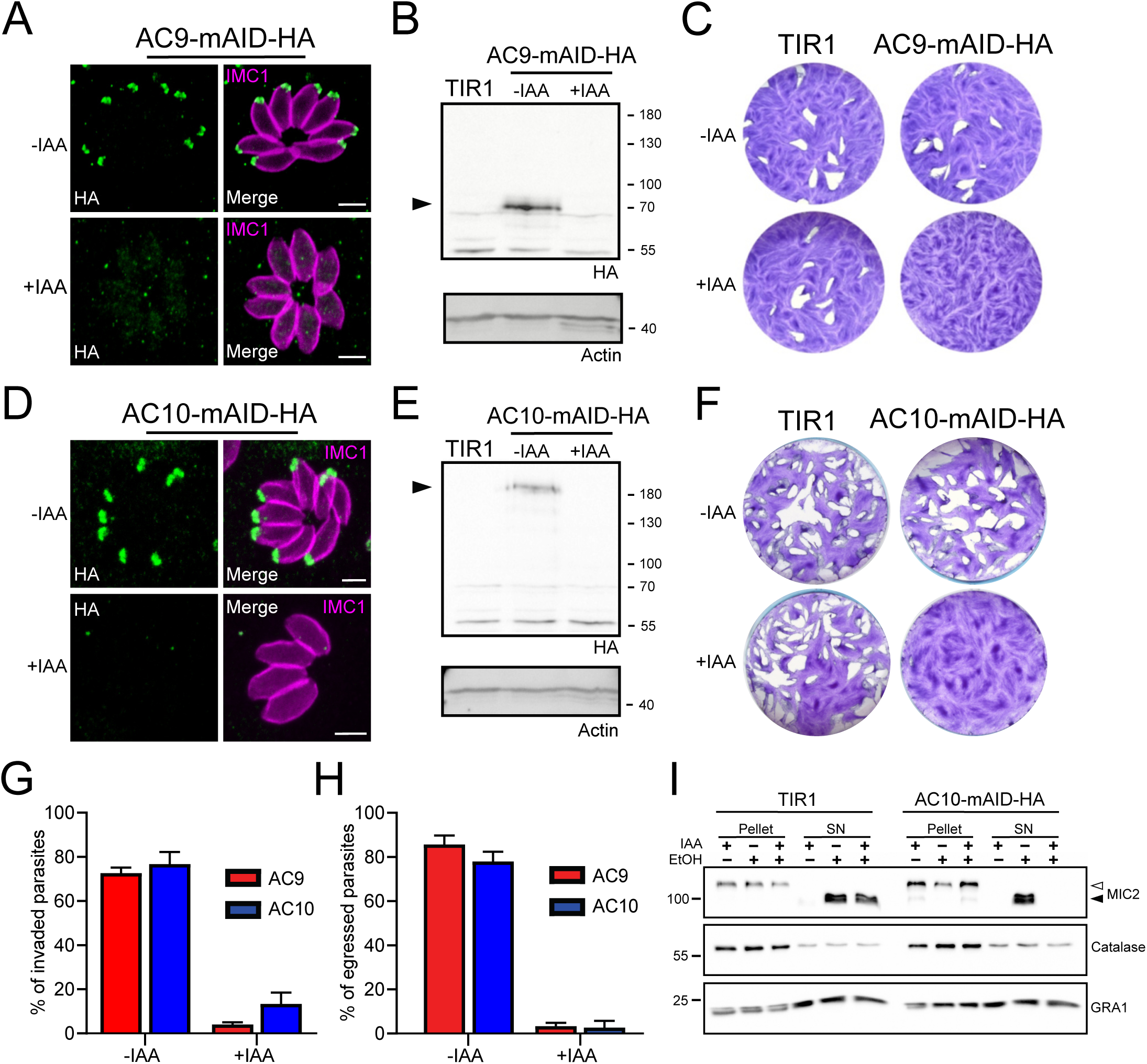
AC9 and AC10 are essential proteins for microneme secretion, invasion and egress. **(A), (B)** and **(C)** AC9-mAID-HA was tightly regulated by addition of IAA and AC9 depleted parasites failed to form lysis plaques after 7 days. TIR1 represents the parental strain and used as control. **(D), (E)** and **(F)** AC10-mAID-HA was rapidly degraded upon addition of IAA and its depletion resulted in no visible plaque formation after 7 days. **(G)** and **(H)** Both invasion and egress were severely impaired in absence of AC9 and AC10. Data represented are mean values ± standard deviation (SD) from three independent biological experiments. **(I)** Microneme secretion was completely abolished in AC10 depleted parasites stimulated by ethanol. Pellets and supernatants (SN) were analyzed using α-MIC2 antibodies for secretion (white arrow: full length MIC2; black arrow: secreted MIC2), α-catalase (CAT) to assess parasites lysis and α-dense granule 1 (GRA1) for constitutive secretion. Scale bars = 2µm.

### Conditional depletion of AC9 or AC10 causes severe morphological defect of the apical complex

In the absence of AC9 or AC10, numerous proteins crucially implicated in motility, invasion and conoid stability were no longer detectable at the apical complex and notably the conoid-associated motor MyoH (20) and the apical polar ring resident protein RNG2 (Fig 5A). Among other apical proteins lost, we found the apical methyltransferase (AKMT) (S5A Fig) (22), the conoid ankyrin-repeat containing protein hub 1 (CPH1) (Fig S5B) and the apical actin nucleator FRM1 (Fig S5C) (15). Conversely, ICMAP1, a MTs binding protein localizing to the intraconoidal microtubules (36) was still detectable but mispositioned (Fig S5D). Remarkably, all the proteins lost in the conoid of the mature parasite (mother) were still present in the forming daughter cells suggesting that these markers are lost during the last step of daughter cell formation.

**Figure 5.**
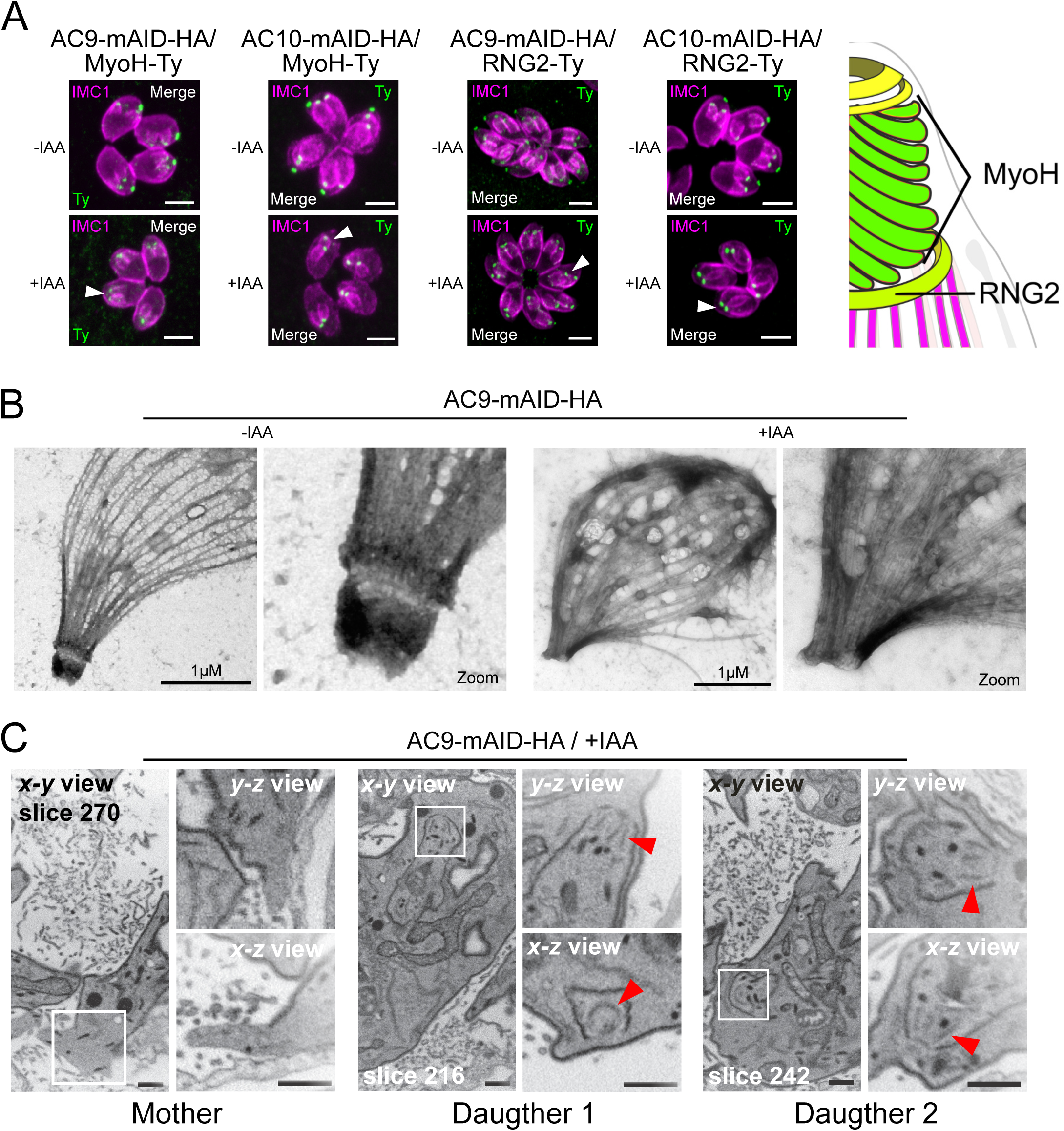
AC9 and AC10 depleted mother parasites lost the APR and the conoid. **(A)** Both conoid-associated MyoH and apical polar ring RNG2 staining were completely lost in mature parasites while still present in forming daughter cell (arrowhead) in absence of AC9 and AC10. **(B)** AC9 depletion caused loss of the conoid and the apical polar ring in agreement with IFA analysis using RNG2 marker. Parasite were grown for 48h −/+ IAA, syringe lysed and processed as described in materials and methods. **(C)** FIB-SEM confirmed that loss of AC9 impacted the conoid and APR exclusively in the mother cell. Red arrows highlight conoids. Scale bars = 2µm.

Extracellular parasites shortly treated with deoxycholate, a detergent which solubilizes the membranes (PM and IMC), offers a better resolution of the parasite cytoskeleton by EM. Strikingly, AC9 treated parasites resulted in the loss of the APR and conoid thus explaining the disappearance of apical markers shown by IFA (Fig 5B). Focus Ion Beam Scanning Electron Microscopy (FIB-SEM) on intracellular parasites compellingly revealed that the conoid is absent in mature parasite while it is still visible in daughter cells thus explaining the loss of apical markers only after completion of parasite division (Fig 5C and S1 Movie). The accumulation of micronemes at the very tip of the parasites was absent in AC9 depleted parasites and, in some cases, the micronemes appeared to be released in the vacuolar space (S5E Fig). Consistent with the sporadic vacuolar staining observed with anti-MIC2 antibodies, some parasites displayed an apical dilatation of the PM into the PV space by EM (S5F Fig). Moreover, rhoptries were still attached to parasite apex but less bundled compared to untreated parasites (S5G Fig).

### The conoid and conoid-associated proteins are lost during division in the absence of AC9

To confirm that loss of the conoid occurred specifically during division, time point experiments were conducted. Importantly, 2 hr of incubation in presence of IAA were sufficient to degrade AC9 to undetectable level by WB in intracellular parasites (S6A Fig) and by IFA on extracellular parasites (S6B Fig). Non-dividing extracellular parasites treated overnight with IAA showed no loss of apical markers as illustrated by staining of MyoH (Fig 6A). In contrast, intracellular parasites treated with IAA at different time points, showed that MyoH disappeared from the conoid only in parasites that had completed at least one cycle of division, i.e. after at least 4h of IAA treatment (Fig 6B). Concordantly, extracellular parasites treated with IAA are not affected in microneme secretion (Fig 6C) whereas treatment on intracellular parasites led to increased microneme secretion defects over time (S6C Fig). The conoid markers analyzed so far are incorporated in the daughter cells since the beginning of the division process, shortly after centrosome duplication. Next, we assessed the fate of APR markers such as KinA and APR1, which are incorporated very early (26) and RNG1 a late marker of division (37). In the absence of AC9 or AC10, APR1 and KinA are lost in mature parasites, but present in daughter cells, respectively (Fig 6D and 6E). We confirmed the late incorporation of RNG1 in daughter cells (Fig 6F and 6G); however, in presence of IAA, RNG1 failed in most of the case, to be properly inserted at the APR shortly before parasites emergence from the mother, resulting in disorganized dots (Fig 6F, 6G and S6D and S6E Fig). Those observations confirmed that the disappearance of the APR, like the conoid, occurs at the final stage of the division process. Interestingly, IFA performed on intracellular parasites fixed earlier post-invasion revealed that the conoid is not lost in a synchronous manner, at least during the first 2 division cycles (Fig S6F). Some apical markers could still be observed at the very late stage giving some hints about a possible explanation of the loss of conoid and APR occurring at the end of division and/or daughter cells emergence when newly formed parasites acquire the PM from the mother (Fig S6F).

**Figure 6.**
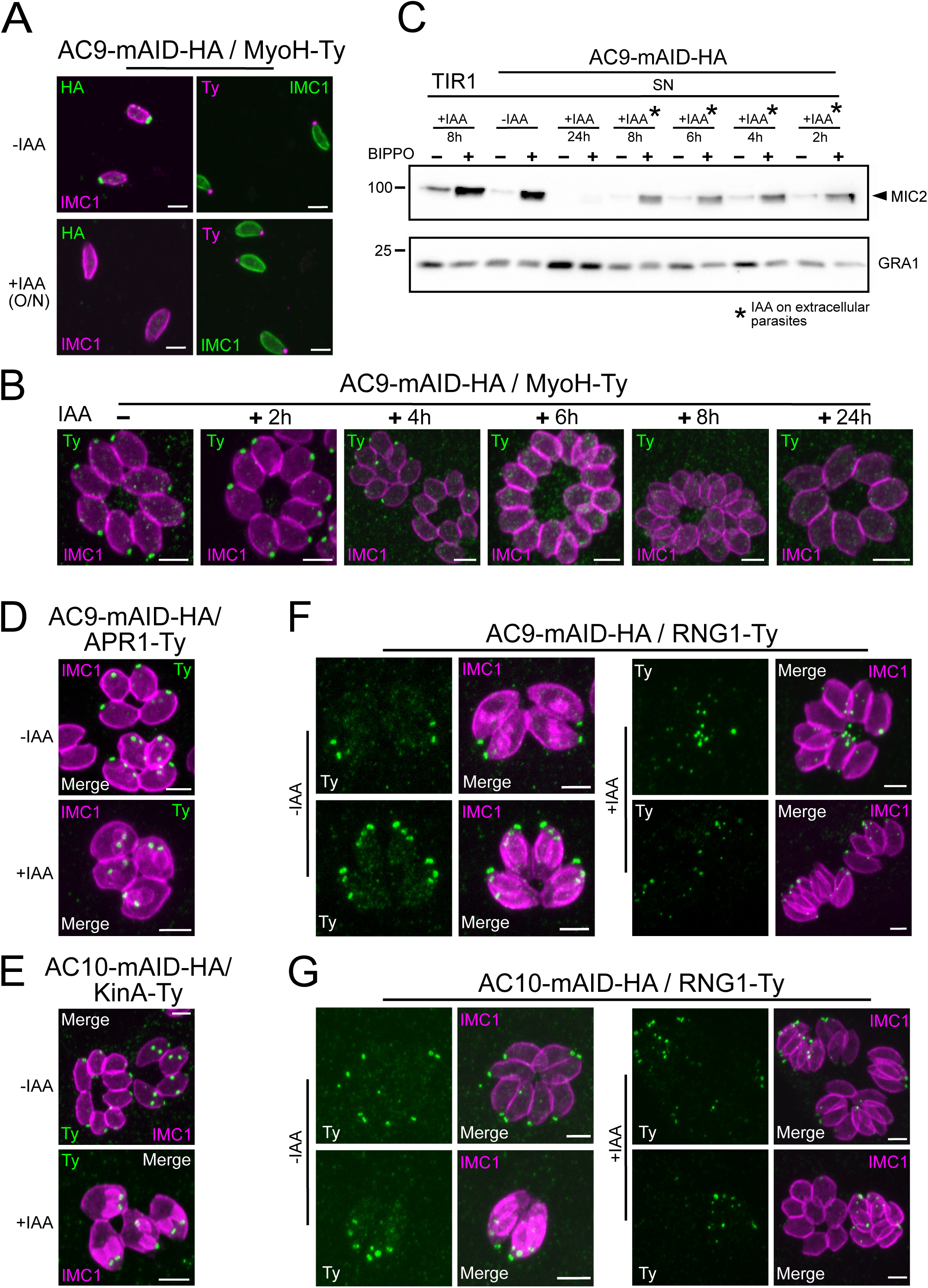
Loss of the APR and conoid occur during the last stages of division. **(A)** IAA overnight treatment of extracellular parasites did not result in MyoH loss from the apical tip of non-dividing mature cell. **(B)** MyoH is lost only during parasite division after more than 2h treatment (i.e. the time to complete at least one division cycle). **(C)** IAA was added at different time point on extracellular parasites showing that microneme secretion is not affected in such conditions. **(D)** and **(E)** APR1 and KinA are lost in mature parasites upon deletion of AC9 and AC10, respectively. **(F)** and **(G)** RNG1 is incorporated very late in daughter cell, way after the alveolin network scaffold stained with IMC1. Upon depletion of both AC9 and AC10, RNG1 failed to be properly anchored at the APR in most parasites. Scale bars = 2µm.

### AC9 and AC10 are required for the assembly of the SPMTs at the apical cap

Given the periodicity of AC9 and AC10 arrangement between the SPMTs and the loss of the conoid and APR upon depletion of two proteins, we wondered if the overall microtubular structure was also impacted. Deoxycholate extraction revealed that in the AC9 and AC10 depleted parasites, the SPMTs were disconnected (Fig 7A and 7B) and only in rare cases the overall structure was maintained despite the obvious absence of APR and conoid (S7A Fig). When applying U-ExM in AC9 and AC10 depleted parasites, loss of APR and conoid were confirmed and the apex appeared enlarged. Remarkably, SPMTs appeared significantly disorganized and primarily disassembled at the apical pole (Fig 7C-F and S7B Fig). Of relevance, the depletion of AC9 resulted in the loss of the preconoidal rings while the peripheral annuli were not affected, when visualized by Centrin2 labeling (Fig 7F).

**Figure 7.**
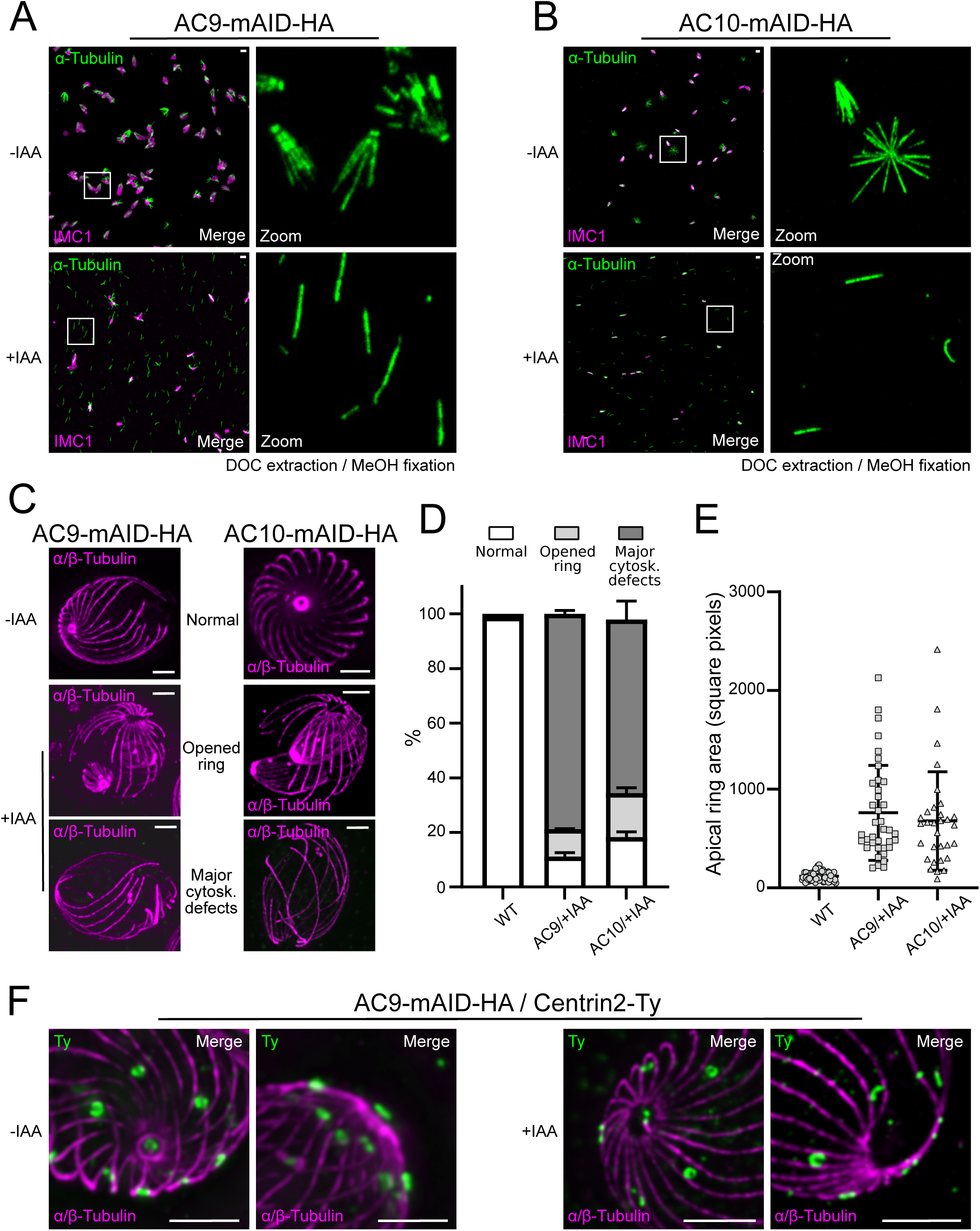
Depletion of AC9 and AC10 caused major cytoskeletal defects. **(A)** and **(B)** Freshly egressed parasites were extracted with DOC on gelatin cover slips. In AC9 and AC10 depleted parasites the microtubular cytoskeleton collapsed and only single microtubules can be visualized. **(C)** U-ExM confirmed that parasites depleted of AC9 and AC10 presented an enlarged apical opening while the microtubular cytoskeleton presented major structural defects. **(D)** and **(E)** Quantification of the cytoskeletal defects and the apical area in absence of AC9 and AC10. **(F)** AC9 depletion caused the loss of pre-conoidal rings while the peripheral annuli were not compromised. Scale bars = 2µm.

**Figure 8.**
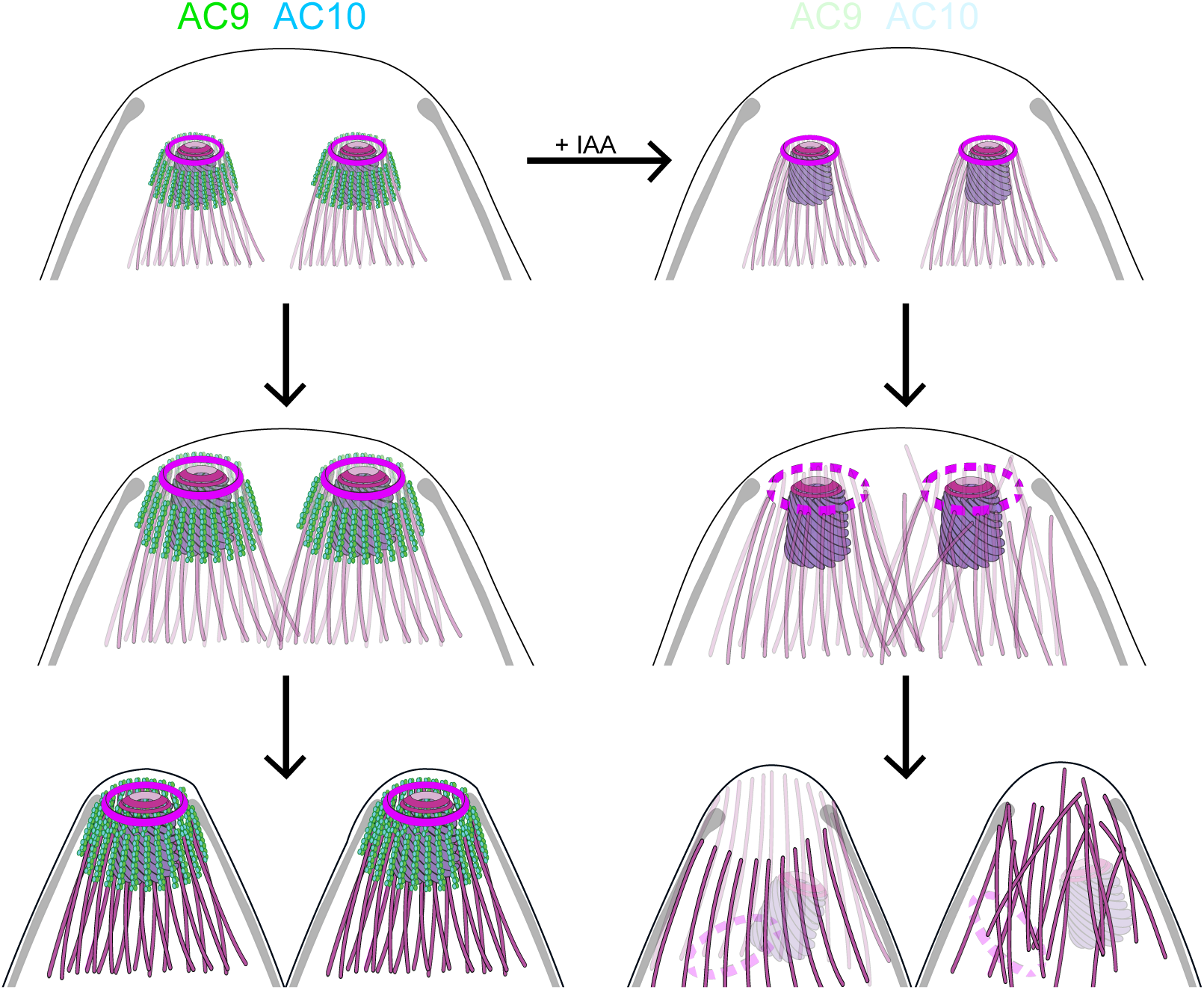
Role of AC9 and AC10 in the stability of the apical complex. Schematic representation of AC9 and AC10 and their role in maintaining the integrity of the structural components of the apical complex during the late stage of parasite division.

## Discussion

In this study we have identified two apical cap proteins implicated in the stability of the APR and anchoring of the conoid to the apical complex of mature parasites. AC10 was identified as the most prominent partner of AC9 via proximity biotinylation of AC9-BirA. Conditional depletion of AC9 or AC10 resulted in a very severe phenotype with parasites impaired in microneme secretion and consequently unable to glide, invade and egress from infected cells. Mature parasites lacking AC9 and AC10 are conoidless and also deprived of APR but replicate normally. In consequence these mutant parasites also fail to assemble the actomyosin system at the conoid that depend on FRM1 and MyoH. In absence of these two factors, no actin filament is produced and delivered to the glideosome, explaining the extreme severity of the phenotype.

### ACs are likely components of the alveolin network conserved in coccidians

Super-resolution microscopy by STED and U-ExM unambiguously established that AC9 and AC10 are not distributed evenly in the apical cap region but organized in longitudinal rows intercalating between the 22 spiraling SPMTs. Fractionation assays indicated that both proteins are poorly soluble consistent with their implication in the alveolin network that spreads between the IMC and the SPMTs. The lack of solubility of AC10 hampered biochemical demonstration that AC9 and AC10 form a complex. However, their perfect colocalization and the fact that AC10 depletion photocopied AC9 and caused its destabilization strongly support their functional and physical association. In addition to AC9 and AC10, AC5 (or TLAP3) was previously suggested to arrange in such longitudinal rows in the apical cap region (38). AC9 and AC10 are the earliest markers of parasite division, prior to ISP1, AC1 and AC5. Like AC9 and AC10, AC8 localizes between SPMTs without being in contact with the APR. In contrast AC2 is coating the SPMTs up to the APR suggesting that it is a microtubule-associated protein (MAPs). It would be informative to assess the precise localization of the other ACs even if the corresponding genes are not predicted to be essential (6) to determine the composition of the alveolin network. Of interest, the first described alveolin protein IMC1 is also arranged in longitudinal rows and not homogeneously distributed around the parasite (1). IMC1 appears to form two rows between each SPMT, again reminiscent of the double rows of IMPs observed by freeze-fracture studies (10). IMC1 is well conserved (39) and harbors a strong negative fitness score however its function in *T. gondii* has not been reported. Of note, AC9 and AC10 are conserved uniquely in the cyst forming coccidian subgroup of Apicomplexa that includes *T. gondii, Neospora caninum, Besnoitia besnoiti* and *Cystoisospora suis*; both proteins are absent in *Eimeria* spp., piroplasmida, haemosporidia and cryptosporidia while other ACs appear to be conserved in other lineage including distantly related gregarines and chromerids. To determine if AC9 and AC10 could potentially interact with other alveolar proteins or MAPs to stabilize the apical complex we examined the localization of ISP, AC2, AC3 and AC8 in parasites depleted in AC9 or AC10 and saw no alteration.

### The conoid and APR are lost as consequence of SPMTs disassembly in the absence of AC9/10

The cascade of events leading to the loss of the conoid and APR involves parasite division. Strikingly, the conoid and APR are present in growing daughter cells and are only lost at the final stage of endodyogeny when the daughter cells emerge and their cytoskeleton confers shape and rigidity to the parasite. The detached conoid appears to be very rapidly degraded because we fail to detect it inside mature parasites even when scrutinizing with FIB-SEM. However, with IFA taken at early stage with parasite that have completed only one or two division cycles, some degrading conoids could be observed with MyoH marker. In bigger vacuoles such staining were not observed because the turnover and recycling of materials in the larger residual body might be faster. Contrastingly, the daughter cells were shown to possess the conoid. In deoxycholate extracted AC9 and AC10 depleted parasites, the SPMTs are formed properly but are not joint together by the APR, which is absent in these mutants. A similar but milder phenotype was also reported in the double knock-out of kinesin A and APR1 with SPMTs partially detached from the parasite apex because of the fragmentation of the APR (26). The strategic position of AC9 and AC10 intercalated between SPTMs suggested a possible implication in maintaining the correct interspacing between the SPMTs. Of note, the stability of SPMTs is maintained by the glideosome-associated protein with multiple-membrane spans family 1 (GAPM1) and other members of the GAPM family (40). Here we have exploited the U-ExM technology to observe the SPMTs at an unprecedented level of resolution using anti-tubulin as well as anti-polyglutamylated (PolyE) antibodies. Of relevance the SPTMs are more glutamylated toward the apical pole of the parasites suggesting that the modification accumulates over time and is absent in the atypical tubulin fiber that form the conoid. The application of U-ExM on the AC9 and AC10 depleted mutant revealed a very striking apical disassembly of the SPMTs. The resulting disordered microtubules could cause the destabilization of the APR, which is present at the top of the SPMTs by presumably serving as MTOC. The dramatic consequences of AC9 and AC10 depletion on SPTMs, APR and conoid occur very late during parasite division (Fig.8); concordantly no major perturbations on the overall shape of the parasite, the IMC, the annuli and the apical positioning of microneme and rhoptries were observed. Taken together the findings lead us to postulate that the loss of the conoid is presumably a consequence of the loss of the APR which could result from the disassembly of the SPMTs at the apical pole of the parasite. In such a model, the AC9/AC10 complex would act as a glue in between SPMTs and establish a functional link between the alveolin network and the SPMTs.

## Abbreviations

AC: apical cap protein
SPMTs: subpellicular microtubules,
IMC: inner membrane complex,
APR: apical polar ring
MTOC: microtubule-organizing center
STED: Stimulated Emission Depletion Microscopy
U-ExM: Ultrastructure expansion microscopy
FIB-SEM: Focus Ion Beam Scanning Electron Microscopy
AiD: Auxin-induced Degron
IAA: Indole-3-acetic acid

## Acknowledgement

We thank the Bioimaging Core Facility, Proteomic Core Facility and Proteins/Peptides Platform (University of Geneva – CMU) for their excellent technical support. We thank the MeBoP students of 2019 for their help in replicating and discussing some of the results presented in this work. We thank Oksana Fiammingo for her excellent technical support and Oscar Vadas for protein expression and purification at the platform of the CMU. We thank Damien Jacot for its critical reading of the manuscript. This work was supported by the European Research Council (ERC) under the European Union’s Horizon 2020 research and innovation program under grant agreement no. 695596 to DSF, the Swiss National Science Foundation (SNSF) 310030_185325 to DSF and PP00P3_187198 to PG, Novartis Foundation for medical-biological Research and ERC ACCENT StG 715289 to PG and the EMBO long term fellowship 284-2019 to EB.

## Author contributions

### Conceptualization

Nicolò Tosetti, Nicolas Dos Santos Pacheco, Dominique Soldati-Favre

### Data curation

Nicolò Tosetti, Nicolas Dos Santos Pacheco, Bohumil Maco, Eloïse Bertiaux and Lorène Bournonville

### Formal analysis

Nicolò Tosetti, Nicolas Dos Santos Pacheco, Bohumil Maco, Eloïse Bertiaux and Lorène Bournonville

### Funding acquisition

Dominique Soldati-Favre, Paul Guichard

### Investigation

Nicolò Tosetti, Nicolas Dos Santos Pacheco, Bohumil Maco, Eloïse Bertiaux and Lorène Bournonville

### Methodology

Nicolò Tosetti, Nicolas Dos Santos Pacheco, Bohumil Maco, Eloïse Bertiaux and Lorène Bournonville

### Project administration

Dominique Soldati-Favre

### Supervision

Dominique Soldati-Favre

### Writing– original draft

Nicolò Tosetti

### Writing– review & editing

Nicolò Tosetti, Nicolas Dos Santos Pacheco, Dominique Soldati-Favre

## Material and methods

### Accession numbers

AC9 (TGGT1_246950) and AC10 (TGGT1_292950).

### Parasite culture

*T. gondii* tachyzoites were grown in human foreskin fibroblasts (HFFs, American Type Culture Collection-CRL 1634) in Dulbecco’s Modified Eagle’s Medium (DMEM, Gibco) supplemented with 5% fetal calf serum (FCS), 2 mM glutamine and 25 μg/ml gentamicin. Depletion of AC9-mAID-HA and AC10-mAID-HA were achieved with 500 μM of auxin (IAA) (30).

### Cloning of DNA constructs

Genomic DNA extractions were performed with the Wizard SV genomic DNA purification kit (Promega). PCRs were performed with Q5 (New England Biolabs) and KOD (Novagen) polymerases. Primers used are listed in the Supplementary File 1. Cloning were performed with E.coli XL-1 Gold chemo-competent bacteria. For endogenous epitope tagging, genomic DNA of the C-terminus of CPH1, RNG2 and ICMAP1 were amplified by PCR, digested with restriction enzymes and cloned into ASP5-3Ty-DHFR (Supplementary File 1) (41). Specific gRNA and KOD PCR were generated using the Q5 site-directed mutagenesis kit (New England Biolabs) on the pSAG1::Cas9-U6::sgUPRT vector (42) for all the other KI constructs listed in Supplementary file 1. AC9-mAID-HA and AC10-mAID-HA were generated by KOD PCR as described in (43); primers for KOD and gRNA are listed in Supplementary file 1.

### Parasite transfection and selection of transgenic parasites

*T. gondii* tachyzoites were transfected by electroporation as previously described (44). Mycophenolic acid (25 mg/mL) and xanthine (50 mg/mL) or pyrimethamine (1 µg/ml) were employed to select resistant parasites carrying the HXGPRT and the DHFR cassette, respectively.

### Antibodies

The antibodies employed are the following: rabbit polyclonal: α-GAP45 (45), α-IMC1 (46), α-ARO (47), α-GAC (15), α-polyglutamate chain (PolyE, IN105) (1:500; AG-25B-0030-C050, AdipoGen) and α-HA (Sigma). Mouse monoclonal: α-ACT (48), α-ISP1 (8), α-SAG1, α-Ty, α-MIC2, ROP2-4 (gifts from J-F Dubremetz, Montpellier) and acetylated α-tubulin (6-11B-1; Santa Cruz Biotechnology). Tubulin antibodies AA344 scFv-S11B (β-tubulin) and AA345 scFv-F2C (α-tubulin). Secondary antibodies Alexa Fluor 405-, Alexa Fluor 488-, Alexa Fluor 594-conjugated goat α-mouse/α-rabbit, were used for IFA. For western blot revelation, secondary peroxidase conjugated goat α-rabbit or mouse antibodies (Sigma) were used.

### Immunofluorescence assay

Parasites were inoculated on HFF cells with cover slips in 24-well plates, grown for 24-30 h, fixed with either 4% PFA/0.05% glutaraldehyde (PFA/GA) or cold methanol, neutralized in 0.1M glycine/PBS for 5 min and processed as previously described (45).

### Confocal microscopy and super-resolution microscopy (STED)

Confocal images were obtained with a Zeiss laser scanning confocal microscope (LSM700 using objective apochromat 63x /1.4 oil) and Leica TCS SP8 STED 3X. Experiments were conducted at the Bioimaging core facility of the Faculty of Medicine at University of Geneva. Confocal images were then processed with ImageJ while STED pictures with LAS X.

### Electron microscopy

For negative staining, extracellular parasite either not treated with IAA or treated with IAA were pelleted in PBS. Conoid protrusion was induced by incubation with 40 µL of BIPPO in PBS for 8 min at 37°C. 4 µL of the sample were applied on glow-discharged 200-mesh Cu electron microscopy grid for 10 min. The excess of the sample was removed by blotting with filter paper and immediately washed on drop of double distilled water. Then, the parasite cytoskeleton was extracted by incubation on two 50 µL droplets of 10 mM deoxycholate in humidified chamber protected from light for 2×5 min. Excess of the detergent was removed by 3 washes on drops of double distilled water and finally the sample was negatively stained with 1% aqueous solution of uranyl acetate for 20 sec and air dried. Electron micrographs of protruded conoid were collected with Tecnai 20 TEM (FEI, Netherland) operated at 80 kV acceleration voltage equipped with side mounted CCD camera (MegaView III, Olympus Imaging Systems) controlled by iTEM software (Olympus Imaging Systems). FIB-SEM samples were prepared as extensively described in (49).

### Invasion assay

Freshly egressed parasites were inoculated on HFF cells with cover slips in 24-well plates, centrifuged for 1 min at 1000 rpm and placed at 37°C for 30 min before fixing for 7 minutes with PFA/GA. Fixed cells were incubated first in 2% BSA/PBS for 30 min, then with α-SAG1 antibodies for 20 min and washed 3 times with PBS. Next cells were fixed with 1% formaldehyde for 7 min, washed once with PBS and permeabilized with 0.2% Triton X-100/PBS for 20 min. Cells were incubated with α-GAP45 antibodies, washed 3 times and incubated with secondary antibodies. 200 parasites were counted for each condition. Data represented as mean values ± standard deviation (SD) (three independent biological experiments).

### Egress assay

*T. gondii* tachyzoites were grown for 30 h on HFF cells with cover slips in 24-well plates. The infected host cells were incubated for 7 min at 37°C with DMEM containing BIPPO prior to fixation with PFA/GA. Immunofluorescence assays were performed as previously described with α-GAP45 antibodies and 200 vacuoles were counted. Data are presented as mean values ± SD (three independent experiments).

### Plaque assay

HFFs were infected with fresh parasites and grown for 7 days before fixation with PFA/GA. After fixation, HFFs were washed with PBS and the host cells monolayer was stained with crystal violet.

### Intracellular growth assay

Parasites were grown for 30 h in 24-well plates prior to fixation with PFA/GA for 10 min. IFA with α-GAP45 antibodies was performed as previously described and the number of parasites per vacuole was counted (200 vacuoles). Data are mean values ± SD (three independent biological experiments).

### Microneme secretion

Freshly egressed parasites were harvested after ∼48h of grown and pellets washed twice in 37 °C pre-warmed intracellular buffer (5 mM NaCl, 142 mM KCl, 1 mM MgCl2, 2mM EGTA, 5.6 mM glucose and 25 mM HEPES, pH 7.2). Parasites were incubated at 37 °C for 15 min in DMEM containing either Ca2+ ionophore, BIPPO or propranolol. Parasite were centrifuged at 1000g for 5 min at 4 °C and supernatants (SN) transferred in different tubes. Pellets were washed once in PBS while supernatants were centrifuged again at higher speed (2000g) for 5 min at 4 °C to clean samples from parasite debris. Pellets and SNs were analyzed by Western blot using α-MIC2, α-catalase (CAT) and α-dense granule 1 (GRA1) antibodies.

### Deoxycholate extraction

Freshly egressed parasites were attached to Poly-L-Lysine-coated coverslips and treated with 10 mM deoxycholate for 20 min at room temperature. Parasites were fixed with methanol for 8 min and proceeded as for IFAs as previously described. Intracellular parasites were treated with IAA for 24h prior to experiment.

### Fractionation assay

Freshly egressed tachyzoites were harvested, washed in PBS and then resuspended in either PBS, PBS/ 1% Triton-X-100, PBS/ 0.1 M Na2CO3, pH 11.5 or SDS. Parasites were lysed by 3 cycle of freeze and thaw in liquid nitrogen and incubated on ice for 30 min. Pellets and supernatants were separated by centrifugation at 4°C for 30 min at 15000 rpm.

### Expression and purification of recombinant AC9 protein from bacteria

Full-length AC9 protein, containing a N-terminal His-10 tag, was subcloned into bacterial expression vector. One and a half liter of BL21 cells expressing His10-AC9 were grown until reaching an OD600 of 0.6. Protein expression was induced by adding 0.5 mM IPTG and leaving the cells to grow for 70 h at 12 °C. Cells were harvested and the pellet was resuspended in 50 ml of lysis buffer (50 mM Tris pH 8, 500 mM NaCl, 15 mM imidazole, 1 mM Triton X-100, 5 % glycerol, 5 mM beta-mercaptoethanol). Cells were lysed by two passages through a French Press at 1000 bar, followed by a brief sonication. Cell debris and membranes were removed by centrifugation at 40’000 g for 35 min at 4 °C. Soluble fraction was passed through a 5 ml His Trap column equilibrated in lysis buffer. After loading, the column was washed with 50 ml of lysis buffer, followed by 30 ml of buffer A (20 mM Tris pH 8, 400 mM NaCl, 15 mM imidazole, 3 mM beta-mercaptoethanol). Protein was eluted with buffer A supplemented with 400 mM imidazole. The eluted protein was concentrated to 1 ml using AMICON 30 MWCO concentrator and applied to a Superdex 200 10/300 increase GL size-exclusion chromatography column equilibrated in buffer C (20 mM Tris pH 7.4, 150 mM NaCl, 3 mM DTT). The pure protein eluted after a volume of 10 ml. Fractions containing pure AC9 were pooled and concentrated to 5 mg/ml before flash freezing in liquid nitrogen.

### Microtubules binding assay

The microtubules binding experiments were performed with the Microtubule Binding Protein Spin-Down Assay Biochem Kit (Cytoskeleton, Inc.) following manufacturer protocol.

### U-ExM

Parasites were centrifuged at 1000 rpm during 5 minutes at 32°C and resuspended in 300µL of PBS 1X. Parasites were sedimented on poly-lysine (A-003-E, SIGMA) coverslips (150µL/coverslip) during 10 minutes at RT. Then parasites were fixed in −20°C methanol during 7 minutes and prepared for Ultrastructural Expansion Microscopy (U-ExM) as previously published (33). Briefly, coverslips were incubated for 5 hours in 2X 0.7 % AA/ 1% FA mix at 37°C prior gelation in APS/ Temed /Monomer solution (19% Sodium Acrylate; 10% AA; 0,1% BIS-AA in PBS 10X) during 1 hour at 37°C. Then denaturation was performed during 1h30 at 95°C (50). Gels were expanded overnight in water and after shrinking in PBS gels were stained 3 hours at 37°C with primary antibodies against HA-tag (1:200), Ty-Tag (1:2.5), PolyE (1:500) and α-tubulin and β-tubulin (1:200). Gel were washed 3 x 10 minutes in PBS-Tween 0.1% prior incubation with secondary antibodies (anti-mouse Alexa 488, anti-mouse Alexa 568, anti-mouse STAR RED anti-rabbit Alexa 488, Anti-mouse Alexa 568, Anti-guinea pig Alexa 568, Anti-guinea pig Alexa 594) (1:400) during 3 hours at 37°C and 3 washes of 10 minutes in PBS-Tween. Overnight a second round of expansion was done in water before imaging. Imaging was performed on a Leica Thunder inverted microscope using 63X 1.4NA oil objective with Small Volume Computational Clearing mode to obtained deconvolved images. 3D stacks were acquired with 0.21 µm z-interval and x,y pixel size of 105nm. Images were analyzed and merged using ImageJ software.

## References

1. Mann T, Beckers C. Characterization of the subpellicular network, a filamentousmembrane skeletal component in the parasite *Toxoplasma gondii*. Molecular & Biochemical Parasitology 2001;115.

2. Gould SB, Tham WH, Cowman AF, McFadden GI, Waller RF. Alveolins, a new family of cortical proteins that define the protist infrakingdom Alveolata. Mol Biol Evol. 2008;25(6):1219–30.

3. Anderson-White BR, Ivey FD, Cheng K, Szatanek T, Lorestani A, Beckers CJ, et al. A family of intermediate filament-like proteins is sequentially assembled into the cytoskeleton of *Toxoplasma gondii*. Cell Microbiol. 2011;13(1):18–31.

4. Frenal K, Dubremetz JF, Lebrun M, Soldati-Favre D. Gliding motility powers invasion and egress in Apicomplexa. Nat Rev Microbiol. 2017;15(11):645–60.

5. Francia ME, Striepen B. Cell division in apicomplexan parasites. Nat Rev Microbiol. 2014;12(2):125–36.

6. Chen AL, Kim EW, Toh JY, Vashisht AA, Rashoff AQ, Van C, et al. Novel components of the Toxoplasma inner membrane complex revealed by BioID. mBio. 2015;6(1):e02357–14.

7. Chen AL, Moon AS, Bell HN, Huang AS, Vashisht AA, Toh JY, et al. Novel insights into the composition and function of the Toxoplasma IMC sutures. Cell Microbiol. 2017;19(4).

8. Beck JR, Rodriguez-Fernandez IA, de Leon JC, Huynh MH, Carruthers VB, Morrissette NS, et al. A novel family of Toxoplasma IMC proteins displays a hierarchical organization and functions in coordinating parasite division. PLoS Pathog. 2010;6(9):e1001094.

9. Hu K, Johnson J, Florens L, Fraunholz M, Suravajjala S, DiLullo C, et al. Cytoskeletal components of an invasion machine--the apical complex of *Toxoplasma gondii*. PLoS Pathog. 2006;2(2):e13.

10. Morrissette N, Murray J, Roos D. Subpellicular microtubules associate with an intramembranous particle lattice in the protozoan parasite *Toxoplasma gondii.* Journal of Cell Science. 1997;110:35–42

11. Hu K, Roos DS, Murray JM. A novel polymer of tubulin forms the conoid of *Toxoplasma gondii.* J Cell Biol. 2002;156(6):1039–50.

12. Morrissette N. Targeting Toxoplasma tubules: tubulin, microtubules, and associated proteins in a human pathogen. Eukaryot Cell. 2015;14(1):2–12.

13. Mondragon R, Frixione E. Ca2+-Dependence of Conoid Extrusion in *Toxoplasma gondii* Tachyzoites. J Eukaryot Microbiol. 1996;43(2):120–7.

14. Lourido S, Shuman J, Zhang C, Shokat KM, Hui R, Sibley LD. Calcium-dependent protein kinase 1 is an essential regulator of exocytosis in Toxoplasma. Nature. 2010;465(7296):359–62.

15. Tosetti N, Dos Santos Pacheco N, Soldati-Favre D, Jacot D. Three F-actin assembly centers regulate organelle inheritance, cell-cell communication and motility in *Toxoplasma gondii.* Elife. 2019;8.

16. Gubbels MJ, Duraisingh MT. Evolution of apicomplexan secretory organelles. Int J Parasitol. 2012;42(12):1071–81.

17. de Leon JC, Scheumann N, Beatty W, Beck JR, Tran JQ, Yau C, et al. A SAS-6-like protein suggests that the Toxoplasma conoid complex evolved from flagellar components. Eukaryot Cell. 2013;12(7):1009–19.

18. Wall RJ, Roques M, Katris NJ, Koreny L, Stanway RR, Brady D, et al. SAS6-like protein in Plasmodium indicates that conoid-associated apical complex proteins persist in invasive stages within the mosquito vector. Sci Rep. 2016;6:28604.

19. Katris NJ, van Dooren GG, McMillan PJ, Hanssen E, Tilley L, Waller RF. The apical complex provides a regulated gateway for secretion of invasion factors in Toxoplasma. PLoS Pathog. 2014;10(4):e1004074.

20. Graindorge A, Frenal K, Jacot D, Salamun J, Marq JB, Soldati-Favre D. The Conoid Associated Motor MyoH Is Indispensable for *Toxoplasma gondii* Entry and Exit from Host Cells. PLoS Pathog. 2016;12(1):e1005388.

21. Long S, Brown KM, Drewry LL, Anthony B, Phan IQH, Sibley LD. Calmodulin-like proteins localized to the conoid regulate motility and cell invasion by *Toxoplasma gondii*. PLoS Pathog. 2017;13(5):e1006379.

22. Heaslip AT, Nishi M, Stein B, Hu K. The motility of a human parasite, *Toxoplasma gondii*, is regulated by a novel lysine methyltransferase. PLoS Pathog. 2011;7(9):e1002201.

23. Jacot D, Tosetti N, Pires I, Stock J, Graindorge A, Hung YF, et al. An Apicomplexan Actin-Binding Protein Serves as a Connector and Lipid Sensor to Coordinate Motility and Invasion. Cell Host Microbe. 2016;20(6):731–43.

24. Nagayasu E, Hwang YC, Liu J, Murray JM, Hu K. Loss of a doublecortin (DCX)-domain protein causes structural defects in a tubulin-based organelle of Toxoplasma gondii and impairs host-cell invasion. Mol Biol Cell. 2017;28(3):411–28.

25. Leung JM, Nagayasu E, Hwang Y-C, Pierce PG, Phan IQ, Stacy R, et al. A doublecortin-domain protein of Toxoplasma and its orthologues bind to and modify the structure and organization of tubulin polymers. BioRxiv. 2019;Preprint.

26. Leung JM, He Y, Zhang F, Hwang YC, Nagayasu E, Liu J, et al. Stability and function of a putative microtubule-organizing center in the human parasite *Toxoplasma gondii*. Mol Biol Cell. 2017;28(10):1361–78.

27. Long S, Anthony B, Drewry LL, Sibley LD. A conserved ankyrin repeat-containing protein regulates conoid stability, motility and cell invasion in *Toxoplasma gondii*. Nat Commun. 2017;8(1):2236.

28. O’Shaughnessy W, Hu X, Beraki T, McDougal M, Reese M. Loss of a conserved MAPK causes catastrophic failure in assembly of a specialized cilium-like structure in *Toxoplasma gondii.* Mol Biol Cell. 2020.

29. Sidik SM, Huet D, Ganesan SM, Huynh MH, Wang T, Nasamu AS, et al. A Genome-wide CRISPR Screen in Toxoplasma Identifies Essential Apicomplexan Genes. Cell. 2016;166(6):1423-35.e12.

30. Brown KM, Long S, Sibley LD. Plasma Membrane Association by N-Acylation Governs PKG Function in *Toxoplasma gondii*. mBio. 2017;8(3).

31. Frenal K, Polonais V, Marq JB, Stratmann R, Limenitakis J, Soldati-Favre D. Functional dissection of the apicomplexan glideosome molecular architecture. Cell Host Microbe. 2010;8(4):343–57.

32. Long S, Brown KM, Sibley LD. CRISPR-mediated Tagging with BirA Allows Proximity Labeling in *Toxoplasma gondii*. Bio Protoc. 2018;8(6).

33. Gambarotto D, Zwettler FU, Le Guennec M, Schmidt-Cernohorska M, Fortun D, Borgers S, et al. Imaging cellular ultrastructures using expansion microscopy (U-ExM). Nat Methods. 2019;16(1):71–4.

34. Howard BL, Harvey KL, Stewart RJ, Azevedo MF, Crabb BS, Jennings IG, et al. Identification of potent phosphodiesterase inhibitors that demonstrate cyclic nucleotide-dependent functions in apicomplexan parasites. ACS Chem Biol. 2015;10(4):1145–54.

35. Carruthers VB, Moreno SN, Sibley LD. Ethanol and acetaldehyde elevate intracellular [Ca2+] and stimulate microneme discharge in *Toxoplasma gondii*. Biochem J. 1999;342(Part 2):379–86.

36. Heaslip AT, Ems-McClung SC, Hu K. TgICMAP1 is a novel microtubule binding protein in *Toxoplasma gondii*. PLoS One. 2009;4(10):e7406.

37. Tran JQ, de Leon JC, Li C, Huynh MH, Beatty W, Morrissette NS. RNG1 is a late marker of the apical polar ring in *Toxoplasma gondii*. Cytoskeleton (Hoboken). 2010;67(9):586–98.

38. Liu J, He Y, Benmerzouga I, Sullivan WJ, Morrissette NS, Murray JM, et al. An ensemble of specifically targeted proteins stabilizes cortical microtubules in the human parasite *Toxoplasma gondii*. Molecular Biology of the Cell. 2016;27(3):549–71.

39. Dubey R, Harrison B, Dangoudoubiyam S, Bandini G, Cheng K, Kosber A, et al. Differential Roles for Inner Membrane Complex Proteins across *Toxoplasma gondii* and *Sarcocystis neurona* Development. mSphere. 2017;2(5).

40. Harding CR, Gow M, Kang JH, Shortt E, Manalis SR, Meissner M, et al. Alveolar proteins stabilize cortical microtubules in *Toxoplasma gondii*. Nat Commun. 2019;10(1):401.

41. Hammoudi PM, Jacot D, Mueller C, Di Cristina M, Dogga SK, Marq JB, et al. Fundamental Roles of the Golgi-Associated Toxoplasma Aspartyl Protease, ASP5, at the Host-Parasite Interface. PLoS Pathog. 2015;11(10):e1005211.

42. Shen B, Brown KM, Lee TD, Sibley LD. Efficient gene disruption in diverse strains of *Toxoplasma gondii* using CRISPR/CAS9. mBio. 2014;5(3):e01114–14.

43. Brown KM, Long S, Sibley LD. Conditional Knockdown of Proteins Using Auxin-inducible Degron (AID) Fusions in *Toxoplasma gondii*. Bio Protoc. 2018;8(4).

44. Soldati D, Boothroyd JC. Transient transfection and expression in the obligate intracellular parasite Toxoplasma gondii. Science. 1993;260(5106):349–52.

45. Plattner F, Yarovinsky F, Romero S, Didry D, Carlier MF, Sher A, et al. Toxoplasma profilin is essential for host cell invasion and TLR11-dependent induction of an interleukin-12 response. Cell Host Microbe. 2008;3(2):77–87.

46. Frenal K, Marq JB, Jacot D, Polonais V, Soldati-Favre D. Plasticity between MyoC- and MyoA-glideosomes: an example of functional compensation in *Toxoplasma gondii* invasion. PLoS Pathog. 2014;10(10):e1004504.

47. Mueller C, Klages N, Jacot D, Santos JM, Cabrera A, Gilberger TW, et al. The Toxoplasma protein ARO mediates the apical positioning of rhoptry organelles, a prerequisite for host cell invasion. Cell Host Microbe. 2013;13(3):289–301.

48. Herm-Götz A, Weiss S, Stratmann R, Fujita-Becker S, Ruff C, Meyhöfer E, et al. *Toxoplasma gondii* myosin A and its light chain: a fast, single-headed, plus-end-directed motor. EMBO J. 2002;21(9):2149–58.

49. Hammoudi PM, Maco B, Dogga SK, Frenal K, Soldati-Favre D. Toxoplasma gondii TFP1 is an essential transporter family protein critical for microneme maturation and exocytosis. Mol Microbiol. 2018.

50. Le Guennec M, Klena N, Gambarotto D, Laporte M, Tassin A, Van den Hoek H, et al. A helical inner scaffold provides a structural basis for centriole cohesion. Science advance. 2020;6(7).

