## supplementray materials for "Two alveolin network proteins are essential for the subpellicular microtubules assembly and conoid anchoring to the apical pole of mature *Toxoplasma gondii*"

#### Supporting information

##### S1 Fig. Proximity biotinylation by AC9

**(A)** Schematic representations of the strategy employed to obtain the AC9-mAID-HA using CRISPR/Cas9 strategy. Red lightning represent Cas9 Induced double-strand brake. **(B)** AC9 localized in mature and is recruited to daughter cells before IMC1. **(C)** Schematic representations of the strategy employed to obtain the AC9-mycBirA strain using CRISPR/Cas9 strategy. **(D)** AC9-mycBirA fusion was correctly localized and upon addition of biotin, enrichment of biotinylated proteins was detected by streptavidin-based fluorophore in both intracellular and extracellular DOC extracted parasites. **(E)** Upon addition of biotin, several biotinylated proteins were detected by western blot. **(F)** Parasites were cultivated for 24h in standard medium supplemented with biotin, harvested, solubilized in cytoskeletal buffer and streptavidin magnetic beads were used to precipitate (IP) biotinylated proteins and sent for mass spectrometry analysis. **(G)** Similar expression profiles of identified proteins by mass spectrometry. Scale bars = 2µm.

##### S2 Fig. No alterations of AC2 and AC8 upon depletion of AC9 and AC10

(A) and (B) AC2 and AC8 are inserted very late during division in the daughter cells after AC9. None of AC2 and AC8 are lost in absence of AC9. (C) Apical cap protein ISP1 was not impacted upon depletion of AC9; however the apical region of one or few parasites per vacuole appeared enlarged (arrow). (D) and (E) AC2 and AC8 were not impacted by depletion of AC10. (F) ISP1 localization is not impacted upon degradation of AC10; similarly to AC9, some parasites presented an enlarged apical opening (arrow). Scale bars = 2µm.

##### S3 Fig. STED and U-ExM revealed unprecedented localization of alveolin network proteins and tubulin modification

**(A)** The alveolin network (stained by IMC1) also appears to be arranged in a periodic pattern (PAF/GLU fixation). Colocalization with tubulin by STED microscopy highlights the localization of IMC1 in between SPMTs (MeOH fixation). **(B)** Colocalization by STED with GAP45 and ISP1 confirmed that AC9 is present in the alveolin network side of the IMC and not in the pellicle. **(C)** Confocal versus U-ExM techniques showing heavy poly-glutamylation of the SPMTs of *T. gondii*. **(D)** Additional images showing the colocalization of AC9 and AC10 at the apical cap by U-ExM. **(E)** Recombinant AC9 was not able to bind microtubules in in vitro assay. MT: microtubule; SN: supernatant; P: pellet. Scale bars = 2μm.

**S4 Fig. AC9 and AC10 did not cause any defect in parasite replication**

**(A)** Intracellular development was not impacted upon depletion of AC9 and AC10. **(B)** AC9 depletion resulted in a complete block of microneme secretion. Extracellular parasites were stimulated with ethanol and BIPPO. Data represented are mean values ± standard deviation (SD) from three independent biological experiments.

**S5 Fig. Destabilization of apical complex components upon depletion of AC9**

**(A)**, **(B)** and **(C)** Apical complex methyltransferase (AKMT), conoid protein hub 1 (CPH1) and apical actin nucleator FRM1 were all lost from the mature tachyzoites but still present in daughter cells (arrowhead). **(D)** ICMAP1 was also lost from the apical tip; however in some parasites per vacuole, ICMAP staining was still present "floating" in the parasite cytoplasm (asterisk). **(E)** Conoidal micronemes (arrowhead) were absent in parasites depleted of AC9 and some vacuoles showed leaking of microneme in the vacuolar space (asterisk). **(F)** By electron microscopy, the conoid was completely lost from mother tachyzoites in AC9 depleted parasites while still present in the growing daughter cells. As showed by IFA, releasing of cytoplasmic material in the vacuolar space was also observed (red asterisk). **(G)** Rhoptries were mildly affected by loss of AC9; they remained attached to the apical tip but appeared to be more dispersed and not compacted together compared to untreated parasites. Scale bars = 2μm.

**S6 Fig. Loss of the APR and conoid just before the emergence of daughter cells**

**(A)** AC9-mAID-HA was rapidly degraded by western blot and **(B)** IFA analysis. **(C)** Intracellular treated parasites lost the ability to secrete microneme in time-dependent manner. **(D)** and **(E)** Additional images showing RNG1 failing to be properly anchored at the APR in most parasites depleted of AC9 and AC10. **(F)** IFA were fixed after approximately 15-16h of IAA treatment resulting in parasite having completed around 1 or 2 division cycles. In such conditions, MyoH staining could be observed in the cytosol or degraded in the last steps of parasite division/daughter cell emergence. Scale bars = 2µm.

**S7 Fig. Structural defects upon depletion of AC9 and AC10**

**(A)** In rare cases shorter treatment with DOC (deoxycholate) did not result in cytoskeleton collapsing but the absence of the apical polar ring and the conoid can be easily observed in IAA treated parasites. **(B)** Additional images of cytoskeletal structural defects in absence of AC9 and AC10. Scale bars = 2µm.

**S1 Table**

Top hits of AC9 BioID identified by mass spec with accession numbers, transcriptomic data, number of peptides detected and CRISPR-Cas9 essentiality score.

**S1 Movie**

FIB-SEM analysis highlighted the specific loss of conoid in the mother cell but not in the growing daughter cells.

### Figure S1

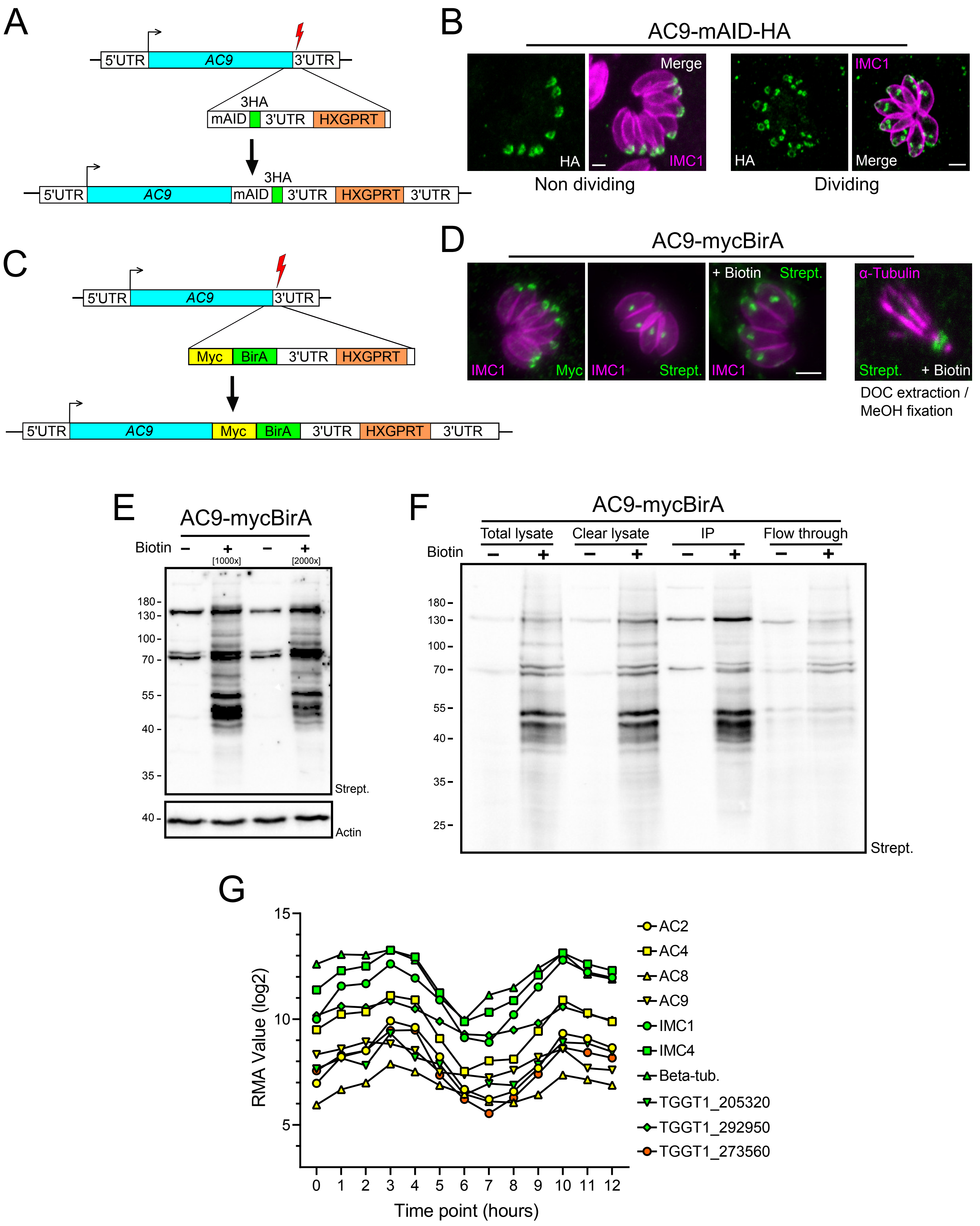

A
5'UTR
AC9
3'UTR
3HA
mAID
HXGPRT
B
AC9-mAID-HA
Merge
HA
IMC1
Non dividing
Dividing
C
Myc
BirA
D
AC9-mycBirA
+ Biotin
Strept.
α-Tubulin
DOC extraction / MeOH fixation
E
Biotin
[1000x]
[2000x]
180
130
100
70
55
40
35
Actin
F
Total lysate
Clear lysate
IP
Flow through
25
G
RMA Value (log2)
Time point (hours)
AC2
AC4
AC8
AC9
IMC1
IMC4
Beta-tub.
TGGT1_205320
TGGT1_292950
TGGT1_273560

### Figure S2

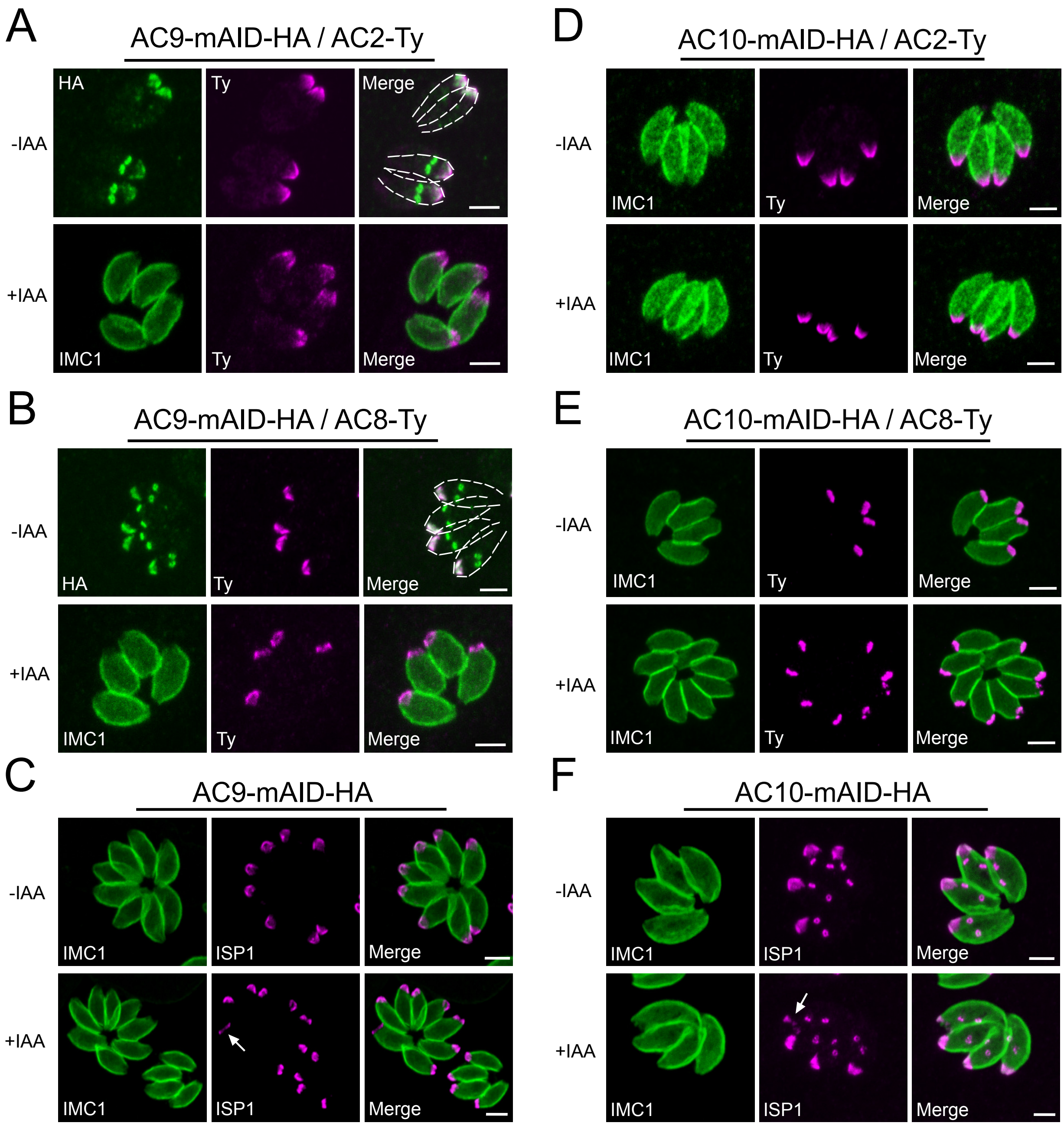

A
AC9-mAID-HA / AC2-Ty
HA
Ty
Merge
-IAA
+IAA
IMC1
Ty
Merge
B
AC9-mAID-HA / AC8-Ty
-IAA
HA
Ty
Merge
+IAA
IMC1
Ty
Merge
C
AC9-mAID-HA
-IAA
IMC1
ISP1
Merge
+IAA
IMC1
ISP1
Merge
D
AC10-mAID-HA / AC2-Ty
-IAA
IMC1
Ty
Merge
+IAA
IMC1
Ty
Merge
E
AC10-mAID-HA / AC8-Ty
-IAA
IMC1
Ty
Merge
+IAA
IMC1
Ty
Merge
F
AC10-mAID-HA
-IAA
IMC1
ISP1
Merge
+IAA
IMC1
ISP1
Merge

### Figure S3

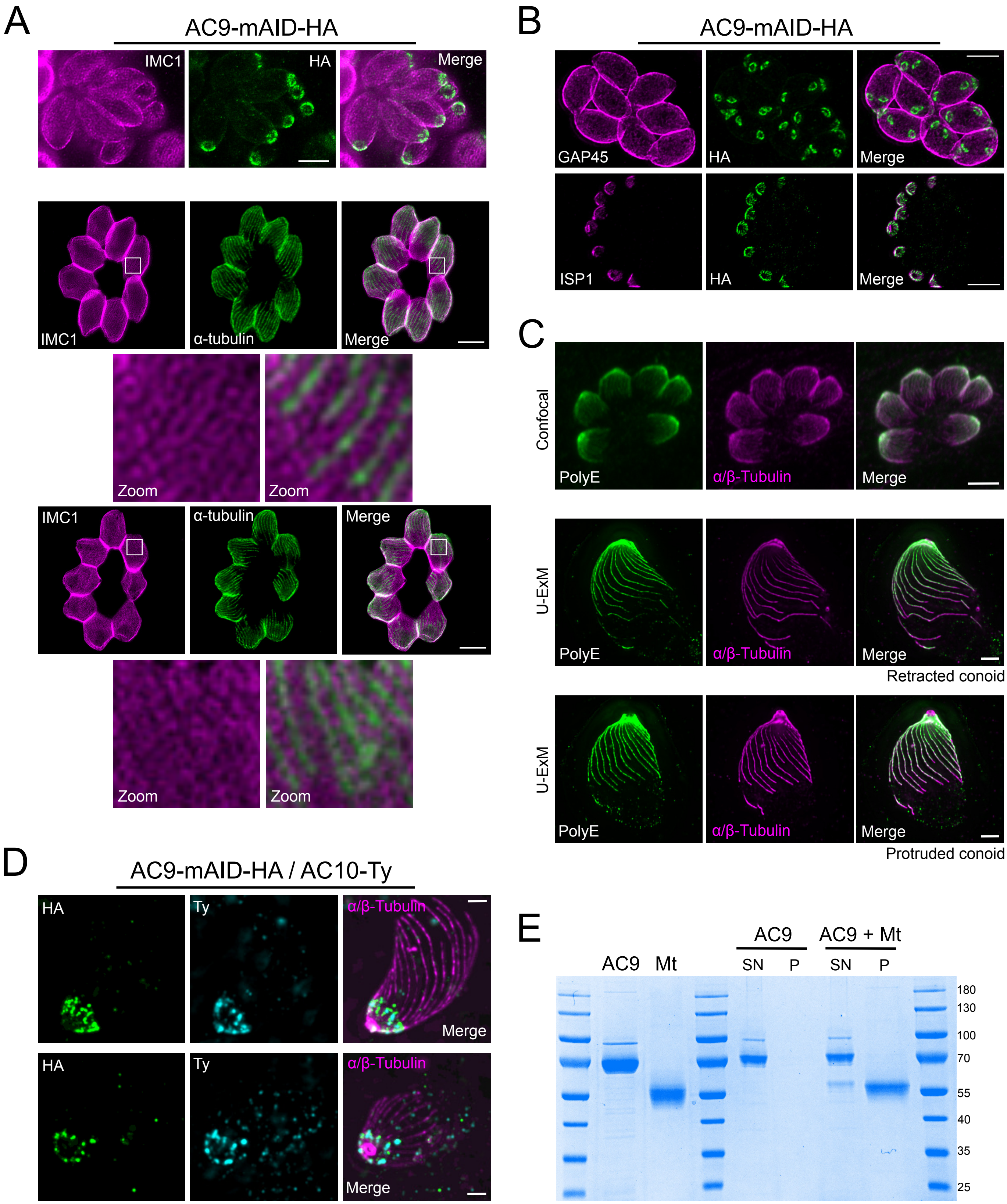

A
AC9-mAID-HA
IMC1
HA
Merge
IMC1
α-tubulin
Merge
Zoom
Zoom
IMC1
α-tubulin
Merge
Zoom
Zoom
B
AC9-mAID-HA
GAP45
HA
Merge
ISP1
HA
Merge
C
Confocal
PolyE
α/β-Tubulin
Merge
U-ExM
PolyE
α/β-Tubulin
Merge
Retracted conoid
U-ExM
PolyE
α/β-Tubulin
Merge
Protruded conoid
D
AC9-mAID-HA / AC10-Ty
HA
Ty
α/β-Tubulin
Merge
HA
Ty
α/β-Tubulin
Merge
E
AC9
Mt
AC9
SN
P
AC9 + Mt
SN
P
180
130
100
70
55
40
35
25

### Figure S4

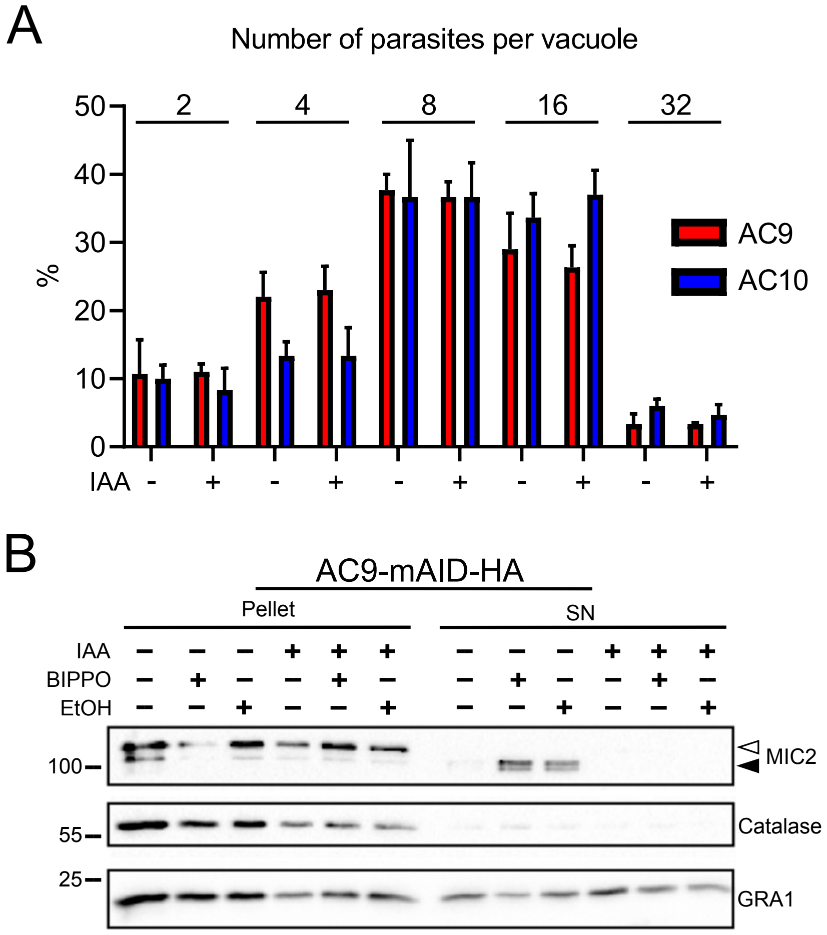
A
Number of parasites per vacuole
2
4
8
16
32
50
40
30
20
10
0
%
AC9
AC10
IAA
-
+
B
AC9-mAID-HA
Pellet
SN
IAA
BIPPO
EtOH
100
55
25
MIC2
Catalase
GRA1

### Figure S5

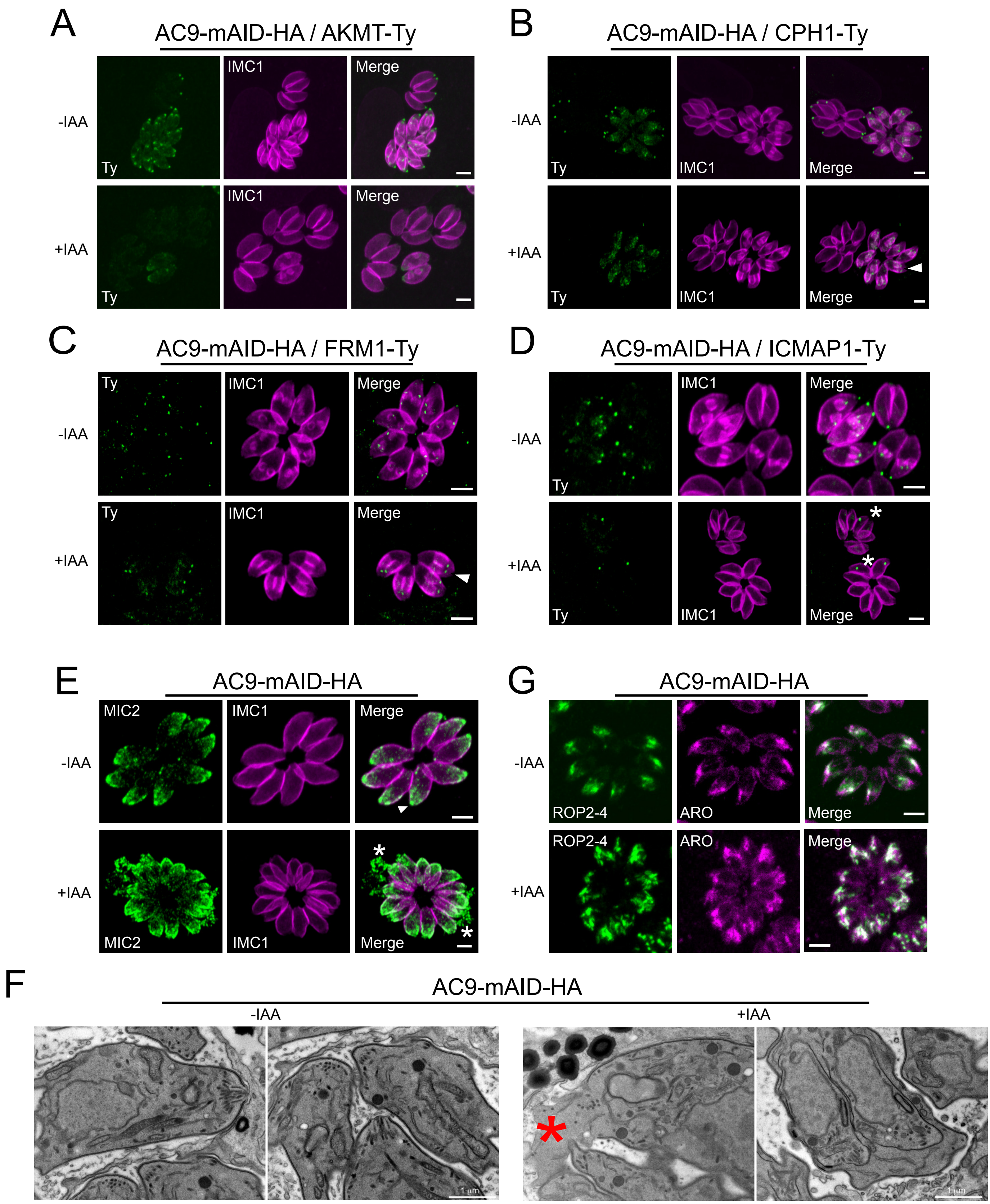

A
AC9-mAID-HA / AKMT-Ty
-IAA
+IAA
Ty
IMC1
Merge
B
AC9-mAID-HA / CPH1-Ty
C
AC9-mAID-HA / FRM1-Ty
D
AC9-mAID-HA / ICMAP1-Ty
E
AC9-mAID-HA
MIC2
G
ROP2-4
ARO
F
1 µm

### Figure S6

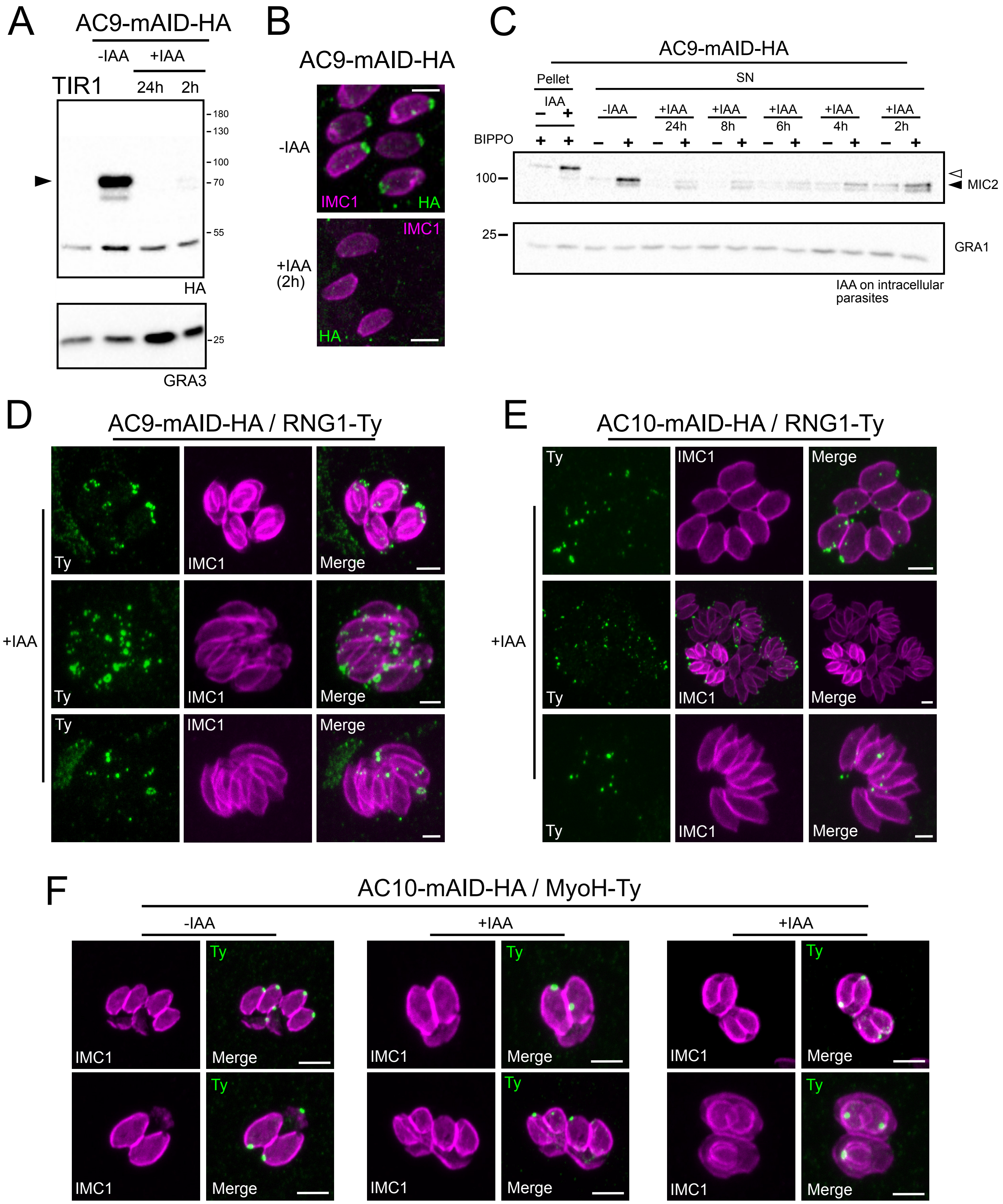
A
AC9-mAID-HA
-IAA
+IAA
TIR1
24h
2h
180
130
100
70
55
HA
25
GRA3
B
AC9-mAID-HA
-IAA
IMC1
HA
+IAA (2h)
IMC1
HA
C
AC9-mAID-HA
Pellet
SN
IAA
-IAA
+IAA 24h
+IAA 8h
+IAA 6h
+IAA 4h
+IAA 2h
BIPPO
100
MIC2
25
GRA1
IAA on intracellular parasites
D
AC9-mAID-HA / RNG1-Ty
+IAA
Ty
IMC1
Merge
E
AC10-mAID-HA / RNG1-Ty
+IAA
Ty
IMC1
Merge
F
AC10-mAID-HA / MyoH-Ty
-IAA
+IAA
+IAA
IMC1
Ty
Merge

### Figure S7

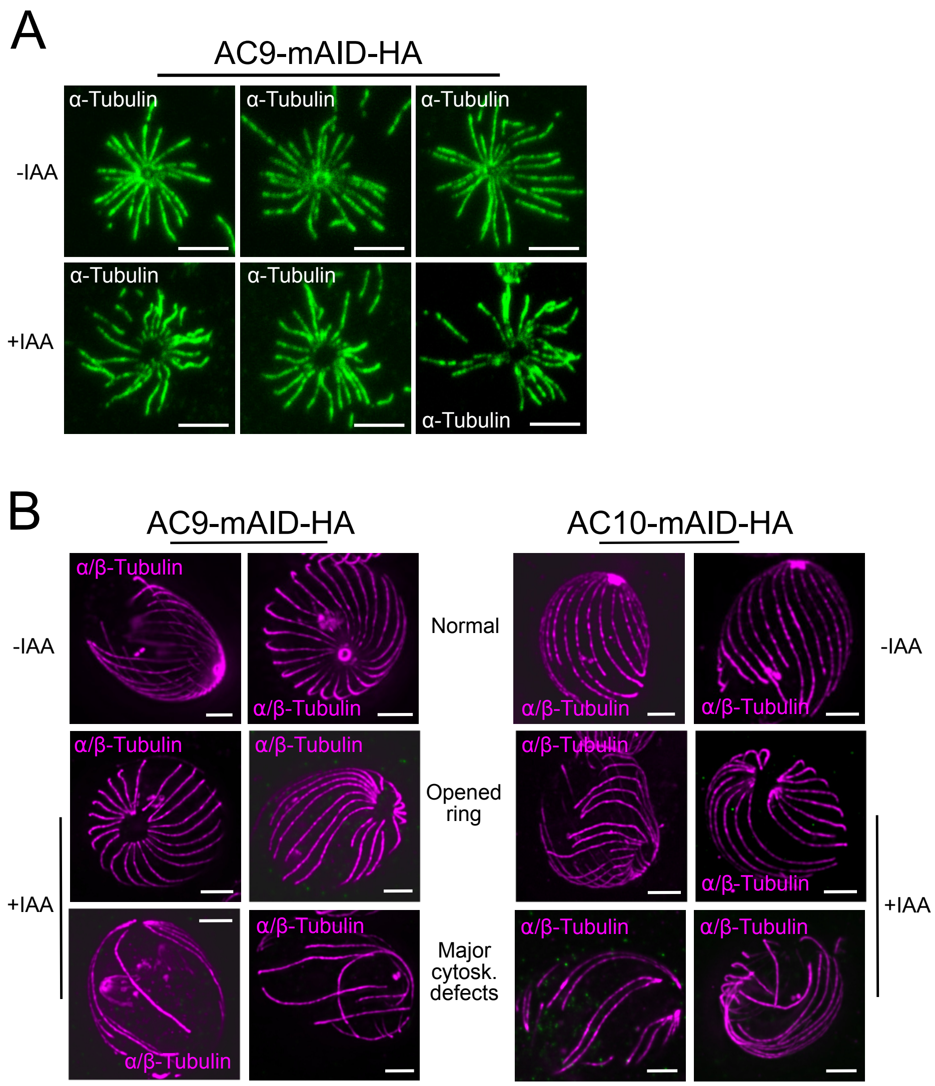
A
AC9-mAID-HA
-IAA
+IAA
α-Tubulin
B
AC9-mAID-HA
AC10-mAID-HA
-IAA
+IAA
α/β-Tubulin
Normal
Opened ring
Major cytosk. defects

### Table S1

| | Accession Number | Transciptomic | IP - Biotin | IP + Biotin | CRISPR's Score |
|---|---|---|---|---|---|
| AC9 | TGGT1_246950 | Late S1 + M | 3 | 14 | -3,88 |
| AC2 | TGGT1_250820 | Late S1 + M | 0 | 27 | 0,5 |
| AC8 | TGGT1_229640 | Late S1 + M | 0 | 27 | -0,1 |
| AC4 | TGGT1_214880 | Late S1 + M | 0 | 4 | 0.01 |
| hypothetical protein | TGGT1_292950 | Late S1 + M | 0 | 55 | -2,58 |
| kinesin heavy chain | TGGT1_273560 | Late S1 + M | 3 | 13 | -0,94 |
| hypothetical protein | TGGT1_205320 | Late S1 + M | 0 | 3 | 0,85 |
| IMC1 | TGGT1_231640 | Late S1 + M | 0 | 13 | -4 |
| IMC4 | TGGT1_231630 | Late S1 + M | 0 | 5 | -4.52 |
| Beta-Tubulin | TGGT1_221620 | Late S1 + M | 0 | 2 | -2,8 |
| hypothetical protein | TGGT1_232340 | ? | 0 | 13 | -0.25 |
| hypothetical protein | TGGT1_213040 | no cycling | 0 | 4 | -4,77 |
| hypothetical protein | TGGT1_209280 | no cycling | 0 | 4 | -0,2 |
| hypothetical protein | TGGT1_214980 | no cycling | 0 | 3 | 0,67 |
| hypothetical protein | TGGT1_261620 | G1 | 0 | 2 | -3,46 |
